# A biologically-grounded cerebellar spiking network model with realistic synaptic transmission captures complex circuit dynamics

**DOI:** 10.64898/2026.05.12.724100

**Authors:** Marialaura De Grazia, Danilo Benozzo, Dimitri Rodarie, Filippo Marchetti, Egidio D’Angelo, Claudia Casellato

## Abstract

Cerebellar neural circuit dynamics rely on a rich repertoire of synaptic and excitability mechanisms, which are thought to determine network computation in physiological and pathological conditions. In this work, we develop and validate a biologically-grounded spiking neural network of the cerebellar cortex, embedding key mechanisms of cellular excitability and synaptic transmission, and assess their impact on signal processing. Neuronal input-output functions, short-term synaptic plasticity, receptor-specific kinetics, and NMDA channel voltage-dependent gating were calibrated against detailed multicompartmental models through automatic tuning procedures. Incorporating these realistic biological properties allowed the network model to simulate key features observed in recordings from acute cerebellar slices. The neuronal discharge and local field potentials elicited by mossy fiber stimulation faithfully reproduced the natural patterns with millisecond precision. Then, selective receptor switch-off revealed the contribution of NMDA, GABA, and AMPA receptors to the frequency-dependent input-output function of the granular layer and Purkinje cells, linking previous empirical findings to specific synaptic mechanisms. This model combines high computational performance with biological realism and offers a computationally efficient framework to investigate neurophysiological phenomena and the neural correlates of behavior in large-scale long-lasting simulations, such as those needed to address the neural underpinnings of learning and of cerebellar pathologies.

## Introduction

The term neural microcircuit generally refers to interconnected groups of neurons within a specific brain region that collectively generate particular functional responses. From this perspective, one of the central aims of neural modeling is to reconstruct and simulate these local circuits, with the goal of capturing the relationships between structure and neuronal dynamics, which in turn generate, or contribute to, complex functional outputs. In this context, we focus on the cerebellar microcircuit. From an anatomical point of view, it is geometrically well defined, organized into three distinct layers, namely the granular layer, the Purkinje cell layer, and the molecular layer. It represents a functional unit of the cerebellum, whose computational principles are involved in multiple domains. Indeed, although the cerebellum has been classically associated with motor control and coordination, it is now well recognized that it also contributes to higher-order functions, including cognitive, emotional, and associative processing [1, 2]. A first attempt toward the development of a general reference model of the mouse cerebellar cortex, hereafter referred to as a *canonical* model [3, 4], was recently provided by a detailed multicompartmental model of this microcircuit [5]. However, beyond detailed multicompartmental microcircuits, spiking neural network (SNN) models made of point neurons are essential for most neurophysiological and technological exploitations and for computational analysis, since they allow massive cost-efficient large-scale simulations. Importantly, the investigation of biophysically grounded computational principles requires cellular excitability, synaptic transmission, and network dynamics to be described through explicit and differentiated mechanisms, with accurately defined spatiotemporal features and robust parameterization. Realistic cerebellar SNNs enable a wide range of applications, including the systematic exploration of model parameter spaces to relate specific mechanisms to network dynamics, their integration into closed-loop systems for sensorimotor and behavioral tasks involving long-term plasticity [6, 7], and derivation of equivalent multi-population cerebellar mean-field description for simulating whole-brain dynamics [8, 9].

As a step toward a unified computational framework for these applications, we developed and validated a realistic SNN of the cerebellar cortex, building on the previously developed multicompartmental representation and proposed here as a canonical SNN model of this microcircuit, in which:

- Point-neuron models were automatically optimized against cell-type-specific multicompartmental models [10–14], exploiting an ad-hoc simplification framework that reproduces the main spiking properties of the different neuronal populations within the network;
- Synaptic dynamics were modeled with receptor-specific components (AMPA, NMDA and GABA), whose parameters were calibrated against synaptic responses generated by multicompartmental model simulations;
- Short-term synaptic plasticity was incorporated to account for dynamic changes in synaptic efficacy;
- Connectivity weights were automatically tuned to match the basal activity regime of the corresponding multicompartmental circuit.

Together, these systematically tuned neuronal and synaptic mechanisms are essential for reproducing the fine temporal filtering, gain modulation, and pathway-specific integration that underlie cerebellar computations [15].

Finally, the cerebellar SNN was validated by quantitatively comparing its responses with ex vivo population recordings obtained from high-density multielectrode arrays (HD-MEAs) in acute cerebellar slices, as well as with additional experimental findings reported in the literature [16–18]. Multiple stimulation patterns delivered through the input fibers were then simulated to characterize nonlinear response dynamics emerging from the interaction among neuronal populations. The same mechanistic framework further enabled the contribution of distinct neuronal populations and synaptic components to be disentangled, allowing selective alterations of synaptic mechanisms, often involved in pathological conditions, to be investigated in a controlled computational setting.

## Materials and Methods

This section describes the methodological workflow leading to the proposed cerebellar SNN, schematically summarized in **Figure 1**.

**Fig 1.**
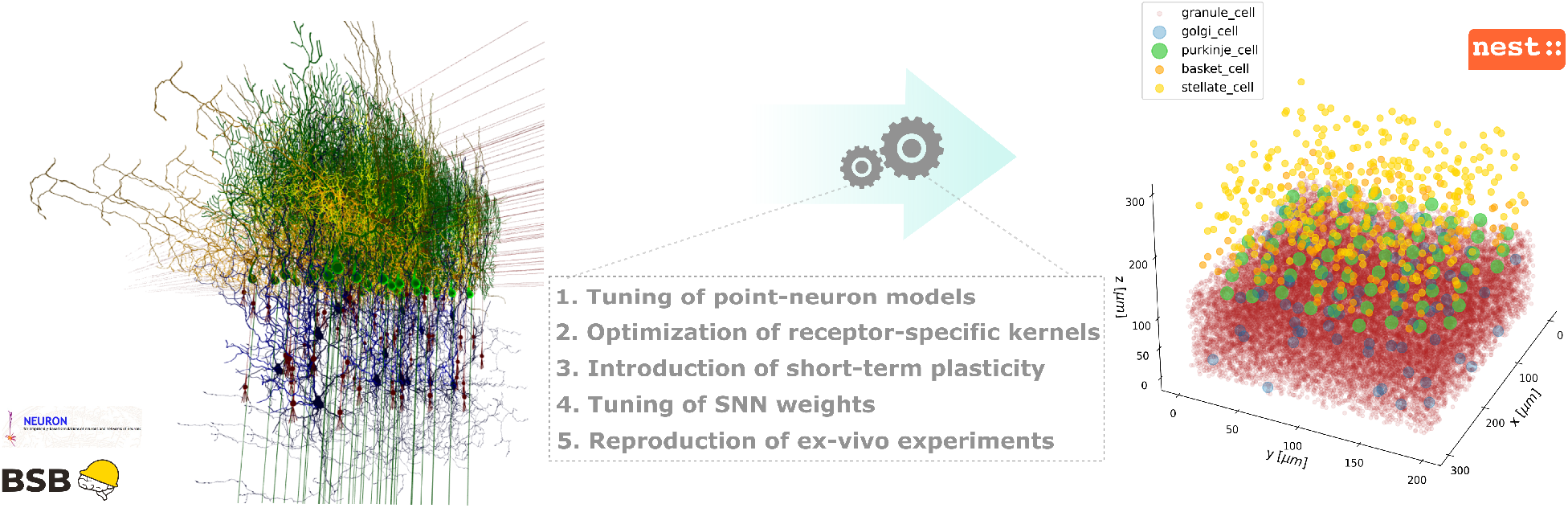
Biophysically-grounded modeling framework for the cerebellar SNN. Cross-scale workflow for calibrating and validating a cerebellar SNN from a detailed multicompartmental reconstruction. Starting from a multicompartmental reconstruction of the cerebellar cortical circuit built with the Brain Scaffold Builder (BSB) and simulated in NEURON [5, 19, 20], a cerebellar SNN model [6, 21] was progressively refined through the tuning of cell-specific point-neuron models, the optimization of receptor-specific synaptic kernels, the incorporation of short-term plasticity mechanisms, and the adjustment of network-level synaptic weights. This cross-scale calibration enabled the SNN to preserve the key dynamical features of the reference multicompartmental model while maintaining computational efficiency. Finally, the ability of the cerebellar SNN to reproduce ex-vivo experimental recordings and literature evidences was evaluated.

Starting from a multicompartmental model of the cerebellar cortex [5], reconstructed with the Brain Scaffold Builder (BSB) toolbox and simulated with the NEURON simulator [19, 20], we significantly extended a previously developed cerebellar SNN framework [6, 21]. Specifically, our aim was to develop a cross-scale procedure yielding a computationally efficient SNN that retains the main multi-component dynamical features of the multicompartmental circuit [5].

### Circuit configuration

The model represents a 300 ×200 ×295 *μm* volume of the mouse cerebellar cortical microcircuit, organized into three main layers:

1. **Granular layer**: the input stage of the cerebellar cortex. Incoming information from mossy fibers (mfs) reaches a dense population of granule cells (GrC), whose axons have an ascending branch before bifurcating into parallel fibers (pf). This layer also contains Golgi cells (GoC), inhibitory interneurons providing feedback inhibition to GrCs. Mfs terminals are organized in specialized structures called glomeruli (glom), which include GrC dendrites and GoC axonal terminals.
2. **Purkinje cell layer**: a thin sheet containing the somata of the Purkinje cells (PC), the main output of the cerebellar cortex projecting to the deep cerebellar nuclei.
3. **Molecular layer**: this layer contains the dendritic trees of PCs and the GrC axons forming parallel fibers (pfs), which establish numerous synaptic contacts with PC dendrites. It also comprises two inhibitory interneuron populations (MLI, molecular layer interneurons), namely stellate (SC) and basket (BC) cells, whose inhibitory action modulates PC activity.

Cell placement and connectivity were implemented within the BSB platform [19], following the configuration proposed by De Schepper et al., 2022 [5]. The main structural features of the resulting network are summarized in **Figure 2**. As shown, the microcircuit exhibits a feed-forward expansion-convergence architecture, in which the dense granule cell population (∼30000 neurons) conveys mossy fiber inputs toward a much smaller PC population (∼70 neurons). Consistently, the connectivity matrices show a marked granule cell to Purkinje cell convergence, with each PC receiving on average ∼1.5 ×10^3^ GrCs inputs. In parallel, granule cells also recruit local inhibitory interneurons, including GoCs, BCs and SCs, which shape and constrain signal propagation within and across layers.

**Fig 2.**
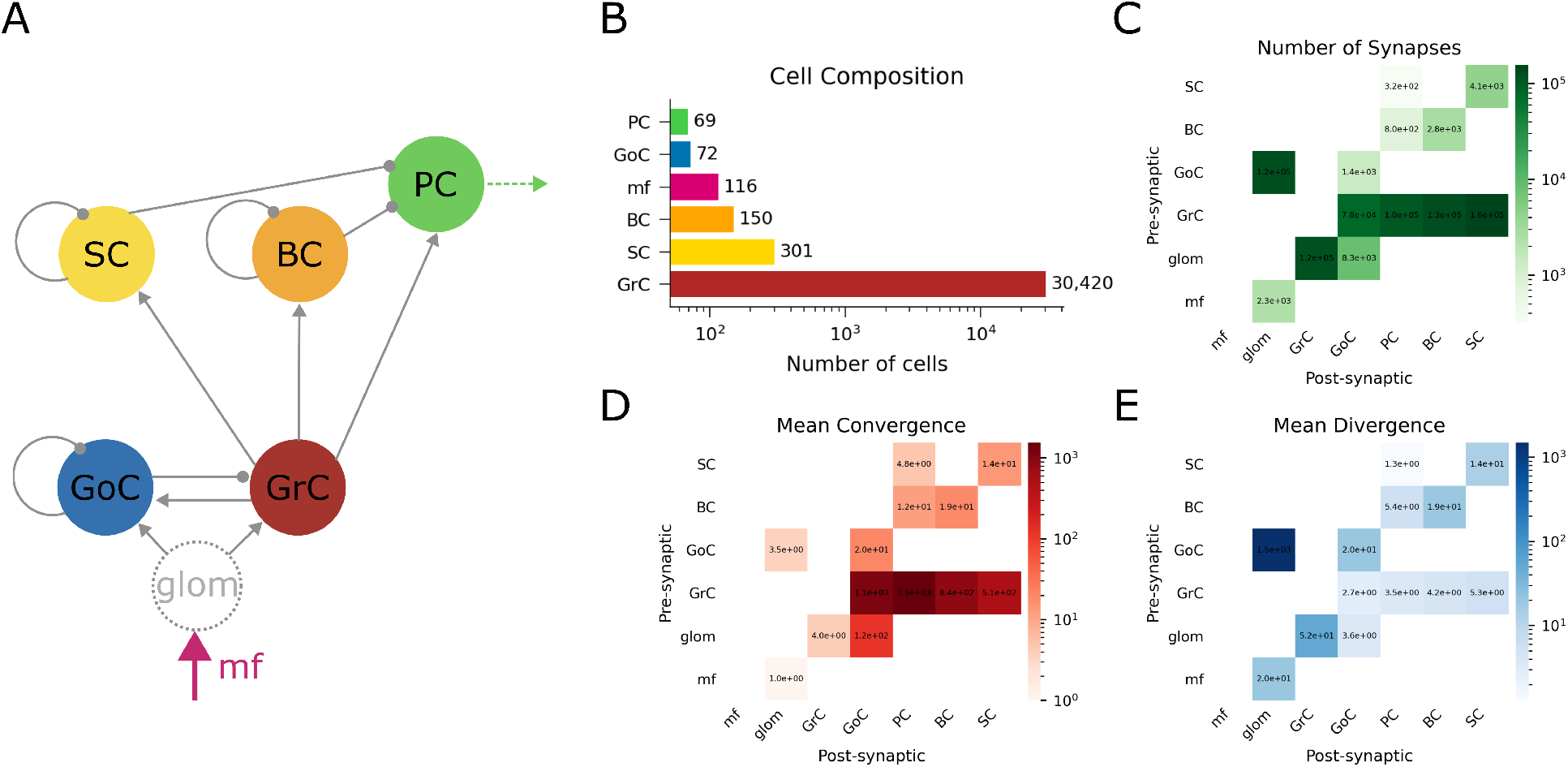
Structural organization of the cerebellar network model. **A**. Schematic representation of the circuit connectivity. **B**. Cellular composition of the network. **C**. Total number of synaptic connections between pre- and post-synaptic populations. **D**. Mean convergence, defined as the average number of pre-synaptic neurons per cell. **E**. Mean divergence, defined as the average number of post-synaptic targets per neuron.

### Single-cell point-neuron model parametrization

Computational modeling often requires linking electrophysiological recordings and biophysically detailed neuronal descriptions to point-neuron formulations suitable for large-scale simulations. Multicompartmental models explicitly represent dendritic morphology and ion-channel dynamics, providing high biophysical realism, but their computational cost limits their direct use in extended networks. Reduced spiking models have therefore been introduced as efficient alternatives [22, 23]. Their predictive value depends on accurate calibration against reference neurons, so that relevant dynamical features are preserved despite the model simplification. Automated optimization strategies have been developed for this purpose, ranging from systematic reductions based on simplified morphologies [24] to data-driven fitting procedures constrained by spike recordings [25, 26].

In this work, cell-specific point-neuron models were calibrated against the corresponding multicompartmental models, apart of mossy fibers (mfs) and glomeruli, which were modeled as simple parrot neurons providing the input to the network. Specifically, the somatic dynamics of neurons were described using the Extended Generalized Leaky Integrate-and-Fire (E-GLIF) formalism [21, 27]. This formalism captures the evolution of the membrane potential and of two intrinsic currents accounting for slow adaptation and fast depolarization, according to:

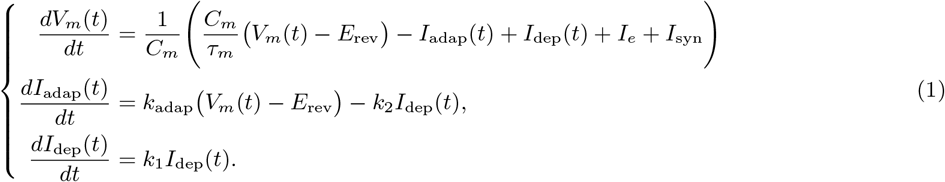

where *I*_syn_ is the total synaptic current, *C*_*m*_ is the membrane capacitance, *τ*_*m*_ is the membrane time constant, *E*_rev_ is the reversal potential, and *I*_*e*_ is the endogenous current. The parameters *k*_adap_ and *k*_2_ regulate adaptation dynamics, whereas *k*_1_ controls the decay of the depolarizing current. When a spike is emitted, the state variables are updated according to:

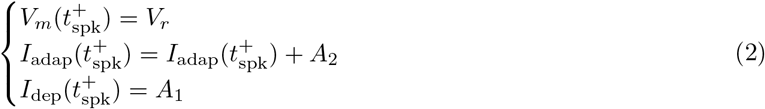

where 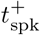 denotes the time instant immediately after spike emission at *t*_spk_, *V*_*r*_ is the reset potential, and *A*_1_ and *A*_2_ control the spike-triggered update of the intrinsic currents.

The automatic tuning procedure was applied to the phenomenological parameters of the E-GLIF model (*A*_1_, *A*_2_, *k*_1_, *k*_2_, *I*_*e*_, *k*_adap_), whereas the remaining biophysical parameters had been previously defined in accordance with the electrophysiological properties of each neuronal population [21, 27]. For each cell type, the goal was to preserve the main electrophysiological features of the corresponding multicompartmental reference model within the reduced point-neuron description. Reference responses were generated through depolarizing and hyperpolarizing current-injection protocols in NEURON [10–14,20]. The same protocols were then simulated with the E-GLIF model, and electrophysiological descriptors were extracted from both model responses. As summarized in Table 1, these descriptors were used to define cell-specific objective (*fitness*) functions, yielding a multi-objective minimization problem that captured the *f* –*I* relationship together with relevant dynamical features such as pacemaking activity, rebound responses, and post-inhibitory pauses, depending on the electrophysiological profile of each cell type.

**Table 1.** Electrophysiological targets used to define the objective functions for the multi-objective optimization of cell-type-specific point-neuron models. For each target, the corresponding fitness quantified the discrepancy between the multicompartmental reference model and the E-GLIF model.

| Electrophysiological feature | Description | Cell types |
| --- | --- | --- |
| **Rheobase** | Minimal current intensity, corresponding to the first current step triggering spike generation. | GrC, GoC, PC, BC, SC |
| **Pacemaking frequency** | Spontaneous autorhythmic firing rate. | GoC, PC, BC, SC |
| **Coefficient of variation (CV)** | Firing variability during autorhythmic activity. | GoC, PC, BC, SC |
| **$f$-$I$ response** | Firing-rate values obtained across input current steps. | GrC, GoC, PC, BC, SC |
| **$f$-$I$ curvature** | Nonlinear profile and saturation behavior of the $f$-$I$ curve. | GrC, GoC, PC, BC, SC |
| **Pause** | Latency to the first spike after stimulation. | PC, BC |
| **Rebound** | Number of spikes after hyperpolarization. | GoC, PC, BC, SC |

The parameter search was implemented in a custom Python workflow based on the DEAP evolutionary framework [28], using NSGA-II algorithm as multi-objective optimizer [29]. This approach was chosen to handle the simultaneous calibration of multiple electrophysiological features over a nonlinear parameter space, a setting in which evolutionary methods have been widely used for neuronal model tuning [30, 31]. NSGA-II returns a Pareto set of non-dominated solutions;therefore, the final parameter set for each cell type was selected using an achievement scalarizing function [32], identifying a balanced trade-off among the different objectives.

### Receptor-specific synaptic transmission

Different receptor types shape synaptic integration over distinct temporal scales and with different effects on neuronal gain, making their differentiation important for mechanistic synaptic modeling [33, 34]. The original E-GLIF model employed generic conductance-based synapses, in which incoming spikes modulated postsynaptic conductance through an alpha-function kernel [27], a common simplified description of synaptic dynamics in spiking neuron models [35, 36]. In the present work, this formulation was extended by introducing receptor-specific synaptic dynamics for AMPA, NMDA, and GABA receptors, in order to account for their distinct temporal and gain characteristics.

Starting from conductance-based synapses with bi-exponential rise–decay dynamics [37], we defined a multi-component formulation in which each receptor type is described by separate fast and slow conductance contributions, enabling a more flexible representation of multi-timescale synaptic dynamics.

Specifically, for each receptor type *X* ∈{AMPA, NMDA, GABA}, the synaptic kernel is described by an auxiliary rising variable *g*_*X,r*_ combined with two decay components, yielding *g*_*X*,fast_(*t*), *g*_*X*,slow_(*t*) from which the total conductance is defined as:

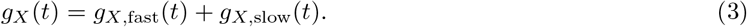

The kernel dynamics is governed by:

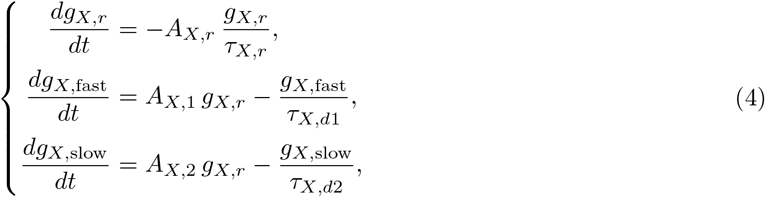

For each receptor type *X*, presynaptic spikes trigger the corresponding synaptic kernel, from which the postsynaptic current is computed as:

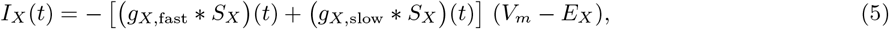

where 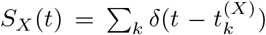 denotes the presynaptic spike train, ∗indicates convolution, *V*_*m*_ is the postsynaptic membrane potential, and *E*_*X*_ is the synaptic reversal potential. The time constants *τ*_*X,r*_, *τ*_*X,d*1_, and *τ*_*X,d*2_ define the characteristic time scales of the kernel dynamics. In particular, *τ*_*X,d*1_ *<τ*_*X,d*2_, so that *g*_*X*,fast_ captures rapid synaptic responses, whereas *g*_*X*,slow_ accounts for slower temporal integration. The parameters *A*_*X,r*_, *A*_*X*,1_, and *A*_*X*,2_ scale the contributions of the different components.

For NMDA synapses, an additional voltage-dependent magnesium (*Mg*^2+^) block was introduced to account for the nonlinear modulation of synaptic current due to *Mg*^2+^ ions. This mechanism reduces NMDA-mediated transmission at hyperpolarized potentials and is progressively relieved upon depolarization. Following the reference multicompartmental models [10], the magnesium block is described by:

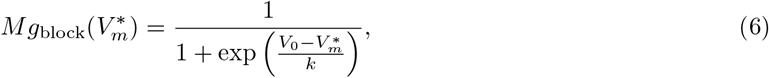

where *V*_0_ = −20 mV and *k* = 13 mV.

Since the present framework employs a LIF neuron model, spike-triggered depolarization was approximated through an auxiliary voltage variable 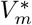, equal to *V*_*m*_ except for a short interval (Δ*t* = 0.5 ms) following each spike, during which it is set to +20 mV. The resulting NMDA current is, therefore:

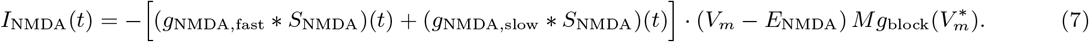

The extended E-GLIF formulation, including the receptor-specific synaptic dynamics described above, was implemented as a dedicated NESTML module, enabling SNN simulations with the new synaptic kernels in the NEST simulator [38–40]. For each connection, and for each receptor component present at that connection, the kernel parameters *g*_init_, *A*_*r*_, *A*_*d*1_, *A*_*d*2_, *τ*_*r*_, *τ*_*d*1_, and *τ*_*d*2_ were calibrated against reference synaptic conductance traces obtained from multicompartmental simulations in NEURON under a voltage-clamp protocol at −40 mV. Here, *g*_init_ represents the initial scaling factor of the kernel.

This calibration was carried out through a standard multi-objective genetic optimization routine proposed by the *pymoo* Python library [41], adopting NSGA-II [29]. Two fitness functions were considered: (1) the mean squared error (MSE) between the conductance traces (from multicompartmental model and from E-GLIF with receptor-specific synaptic kernels), under a synaptic single-spike protocol, and (2) the percentage error on peak amplitude.

The final parameter set was selected from the Pareto front as the solution achieving a peak-amplitude error below 1%and the minimum MSE.

### Short-term synaptic dynamics

Short-term plasticity (STP) dynamically modulates synaptic efficacy over short time scales, thereby regulating excitatory and inhibitory information flow in cerebellar circuits [42]. In the cerebellar SNN, STP was modeled using the Tsodyks–Markram formalism [43], with synaptic parameters assigned according to the corresponding connection type. This connection-specific implementation follows the rationale of recent spiking microcircuit models incorporating cell-type-specific STP to investigate its impact on circuit dynamics [44]. For each connection type, the parameters *U, τ*_fac_, *τ*_rec_, and *τ*_psc_ were adopted from Masoli et al., 2022 [42] (see Table S1 for the complete parameter set).

A key methodological step in the implementation of the STP model was to preserve consistency between static and plastic network simulations [44]. To achieve this, a synaptic weight rescaling factor relative to the corresponding static value was computed so that, for each receptor type and connection, the area under the synaptic conductance curve matched that of the static model under low-frequency stimulation (5 Hz), assuming that STP effects were negligible in this regime. The resulting STP mechanisms were then assessed using single-cell stimulation protocols with presynaptic spike trains at different input frequencies.

### Electrophysiological data

To validate the cerebellar SNN against experimental recordings, we used internal datasets of local field potential (LFP) recordings of the granular layer, obtained from acute parasagittal cerebellar slices of healthy mice. The slices were placed on a high-density multielectrode array (HD-MEA) platform (BioCAM X, 3Brain AG, Wädenswil, Switzerland), consisting of a 64×64 matrix of microelectrodes. Neuronal activity was acquired using BrainWave X software (3Brain AG) at a sampling rate of 17,840.7 Hz per electrode.

Spontaneous activity was first recorded. Then, a single-pulse stimulation protocol was applied, consisting in the delivery of a single 50 *μ*A current pulse through a bipolar stimulating electrode. For each slice, the protocol was repeated 30 times at a frequency of 0.1 Hz.

The HD-MEA recordings were analyzed to characterize the network responses evoked by mossy fiber stimulation in the cerebellar granular layer, which elicited LFP responses characterized by the typical N1–N2A–N2B–P2 complex. Within this waveform, N1 reflects the presynaptic volley, N2A and N2B are associated with GrCs synaptic activation, whereas P2 represents the return current from the molecular layer [16, 17, 45]. Since one of the main aims of the analysis was to compare GrCs synaptic responses between experiments and simulations, N2A and N2B components were considered.

To analyze these components, stimulus-related artifacts were first identified in the raw traces using peak detection, allowing the extraction of 30 individual trials. For each trial, a time window including 200 ms of pre-stimulus baseline and 500 ms of post-stimulus activity was selected. Signals were low-pass filtered with a cutoff frequency of 1000 Hz, and stimulation artifacts were subsequently removed by applying a blanking window from –4.5 ms to +1.5 ms around each detected artifact peak.

For each trial, baseline correction was performed by subtracting the mean of the pre-stimulus segment. Channel selection was carried out based on the signal-to-noise ratio, computed as the peak-to-peak amplitude divided by the Root Mean Square of the baseline across all trials, and on a consistency metric, defined as the mean of the correlation coefficients between individual trial traces and the average trace across all trials. Channels with insufficient signal-to-noise ratio (<5) or low trial-to-trial consistency (*<* 0.25) were excluded from further analysis. For the valid channels, trial-wise peak detection was performed to extract the amplitudes and latencies of the N2A and N2B negative peaks. The N2A peak was searched within the interval [0, 3.0] ms, whereas the N2B peak within [3.0, 8.0] ms.

This analysis can provide quantitative measures of granular layer activation, enabling a direct comparison between simulated and experimental responses, while ensuring a robust and reproducible evaluation framework.

### Cerebellar SNN simulations

Based on the geometrical reconstruction of the network generated with the BSB framework [5, 19], we used the spatial distribution of neurons and their connectivity to simulate the dynamics of the cerebellar microcircuit through the BSB–NEST simulation interface [38].

#### Synaptic weights tuning

Prior to the main simulation experiments, we performed a SNN weight-tuning procedure to reproduce population firing rates comparable to those of the multicompartmental network model [5], under a basal activity protocol, representing the in vivo-like spontaneous dynamics of the network in the absence of specific sensory inputs. The only afferent drive was provided, for 5 s, through mossy fibers by a 4 Hz Poisson process, as background activity [46].

In particular, the multi-objective optimization pipeline previously adopted for single-cell point-neuron parametrization was extended to the network level for setting the synaptic weights of the cerebellar SNN. The fitness functions were defined as the mean population firing rate and its standard deviation for each neuronal population, as reported in [5].

This further tuning step was required because in point-neuron models, synapses are effectively collapsed onto the soma, while, in the multicompartmental neuronal description, dendritic filtering effects are explicitly represented [47]. Therefore, the optimized SNN weights provide an effective phenomenological compensation for the loss of dendritic attenuation when moving from spatially extended neurons to simplified point-neuron representations.

Synapses targeting granule cells (GrCs) were excluded from this correction, as their limited dendritic extent is expected to minimize dendritic filtering effects. Likewise, standard unitary weights were retained for mossy fibers and glomerular relay elements, which were modeled as parrot neurons.

For all remaining connection types, a single effective synaptic weight was optimized. In excitatory connections including both AMPA and NMDA components, the same connection-level weight was assigned to the two receptor components, since they are co-activated by the same presynaptic events and share the same STP parametrization, while receptor-specific kinetics and conductance profiles remained defined by the calibrated synaptic kernels.

#### Network simulations

The tuned network was then simulated with several stimulation protocols, to investigate the dynamical behavior of the model:

- *Single-pulse stimulation protocol* : to reproduce the standard single-pulse mossy fiber stimulation commonly adopted in ex vivo recordings from acute cerebellar slices, we first simulated 1 s of spontaneous activity, after which a synchronous spike was delivered across the entire mossy fiber bundle, without background input. This condition was designed to mimic the experimental setting, in which the granular layer is typically quiescent immediately before stimulation, thus allowing the isolated evoked response of the cerebellar cortex to be examined.
- *Mf stimulation at different frequencies*: to characterize short-term plasticity and frequency-dependent network dynamics, the mossy fiber bundle was stimulated with five-spike trains delivered at frequencies *f* ∈{1, 6, 10, 20, 50, 100, 200, 250, 500} Hz, matching those adopted in previous experimental studies [16–18].

Spike trains were recorded from all neurons in the network.

The described protocols were also repeated by modulating NMDA and GABA receptor strengths, to assess their relative roles to network dynamics. Indeed, in addition to complete receptor blockade, receptor deficit and excess conditions were explored in a continuum, by selectively scaling the receptor-specific synaptic components.

For the evaluation and validation of network performance, we focused on reproducing two main readouts: Purkinje cell spiking activity, validated against experimental evidence reported in the literature, and granular layer LFPs obtained from MEA recordings of cerebellar acute slices. While Purkinje cell spiking activity can be directly recorded within the simulation framework using standard spike detectors, in-silico granular layer LFPs require the definition of a suitable proxy signal. In point-neuron models, extracellular potentials cannot be directly reconstructed, since LFPs arise from spatially distributed transmembrane currents [48], which are not explicitly represented in simplified neuronal descriptions. Nevertheless, combinations of synaptic currents have been shown to provide reliable proxies of LFP temporal dynamics in spiking neural networks [49]. In the cerebellar granular layer, the use of such a proxy is further supported by the anisotropic organization of the microcircuit: GrCs are not recurrently connected to each other, and their axons follow a stereotyped geometry, with ascending branches crossing the granular layer before bifurcating into parallel fibers in the molecular layer. Together with the electrotonically compact structure of granule cells [50], this spatially ordered arrangement makes the aggregate synaptic current dynamics a plausible approximation of the local LFP temporal profile.

On this basis, we defined a virtual recording grid composed of spherical sampling volumes, each representing an LFP channel and collecting the synaptic activity of the GrCs within its volume. The centers of the spheres were positioned at mid-depth of the granular layer (*z* = 65 *μ*m) and arranged on the *x*–*y* plane over the ranges [50, 300] *μ*m and [50, 200] *μ*m, respectively, with a spatial step of 50 *μ*m. Each sphere had radius *r* = 10 *μ*m, chosen to contain on average about 15–20 GrCs, in agreement with experimental estimates for HD-MEA channels [16, 17]. This configuration resulted in a total of 15 virtual recording channels. For each channel *c*, the LFP proxy signal was computed as

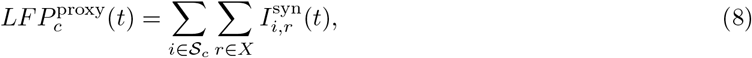

where 𝒮_*c*_ denotes the set of GrCs contained in the *c*-th sphere, *X* is the set of receptor-mediated synaptic components considered in the model (here, AMPA, NMDA and GABA), and 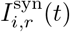 is the synaptic current mediated by receptor type *r* in the *i*-th GrC.

The resulting signal should therefore be interpreted as a simplified and spatially localized proxy of experimentally measured LFPs, designed to capture the main synaptic-current dynamics of the local response, although not intended as a full biophysical reconstruction of the extracellular potential.

To enable a direct quantitative comparison with the experimental dataset, the peak detection procedure applied to the ex vivo recordings was also used for the simulated LFP proxy signals. The analysis was restricted to the N2A and N2B components, as these reflect the synaptic part of the response and are therefore the components most directly represented by the adopted proxy.

## Results

### Cell-specific and synapse-specific parametrization

The optimization steps at the single-cell and single-synapse levels were designed to reproduce the main electrophysiological and synaptic responses by minimizing the mismatch between the multicompartmental models and the corresponding point-neuron representations.

The single-cell tuning procedure generated E-GLIF models able to reproduce the main cell-specific spike pattern properties of the corresponding multicompartmental references (see Table 2). The resulting fitness errors were below 12%for all objective functions (see Table 3), indicating that the reduced point-neuron formulation preserved the relevant electrophysiological features targeted by the multi-objective optimization.

**Table 2.** Optimized E-GLIF phenomenological parameters for each cerebellar cell type. The reported values define the cell-specific intrinsic dynamics of the point-neuron models after the automatic tuning procedure.

| Cell type | $k_{\text{adap}}$ | $k_2$<br>[ms<sup>-1</sup>] | $A_2$<br>[pA] | $k_1$<br>[ms<sup>-1</sup>] | $A_1$<br>[pA] | $I_E$<br>[pA] |
| --- | --- | --- | --- | --- | --- | --- |
| GrC | 0.0675 | 0.0410 | 4.20 | 0.160 | 20.7 | 4.91 |
| GoC | 0.110 | 0.0230 | 94.2 | 0.0680 | 345 | 24.5 |
| PC | 0.481 | 0.0350 | 4.96 | 0.288 | 1.95e3 | 159 |
| BC | 0.175 | 0.114 | 128 | 0.161 | 192 | 5.23 |
| SC | 0.175 | 0.112 | 164 | 0.157 | 209 | 11.1 |

**Table 3.** Errors associated with the fitness functions used for the optimization of each cerebellar cell type. Each term quantifies the discrepancy between multicompartmental and E-GLIF responses for the corresponding electrophysiological target, and is reported as a percentage.

| Cell type | Rheobase<br>error [%] | Pacemaking<br>Frequency<br>error [%] | Coefficient of<br>Variation (CV)<br>error [%] | $f$-$I$ Response<br>error [%] | $f$-$I$ Curvature<br>error [%] | Pause<br>error [%] | Rebound<br>error [%] |
| --- | --- | --- | --- | --- | --- | --- | --- |
| GrC | 0.0 | — | — | 2.8 | 5.6 | — | — |
| GoC | 0.0 | 9.5 | 5.9 | 4.1 | 10.4 | — | 10.0 |
| PC | 0.0 | 5.0 | 6.8 | 8.1 | 11.8 | 5.5 | 12.0 |
| BC | 0.0 | 7.7 | 6.2 | 6.9 | 7.6 | 4.2 | 8.3 |
| SC | 0.0 | 7.7 | 6.7 | 4.7 | 8.6 | — | 0.0 |

**Figure 3** illustrates the outcome of this procedure in detail for the Purkinje cell model. The comparison between the multicompartmental target and the E-GLIF model shows a close agreement in the *f*–*I* relationship, while the individual fitness errors quantify the residual mismatch associated with each electrophysiological descriptor. Representative voltage traces further show that the reduced model captured the main firing regimes of the reference cell model, including spontaneous tonic discharge, increased firing under depolarizing current injection, and firing suppression during hyperpolarization, as illustrated for +300 pA and −300 pA current steps.

**Fig 3.**
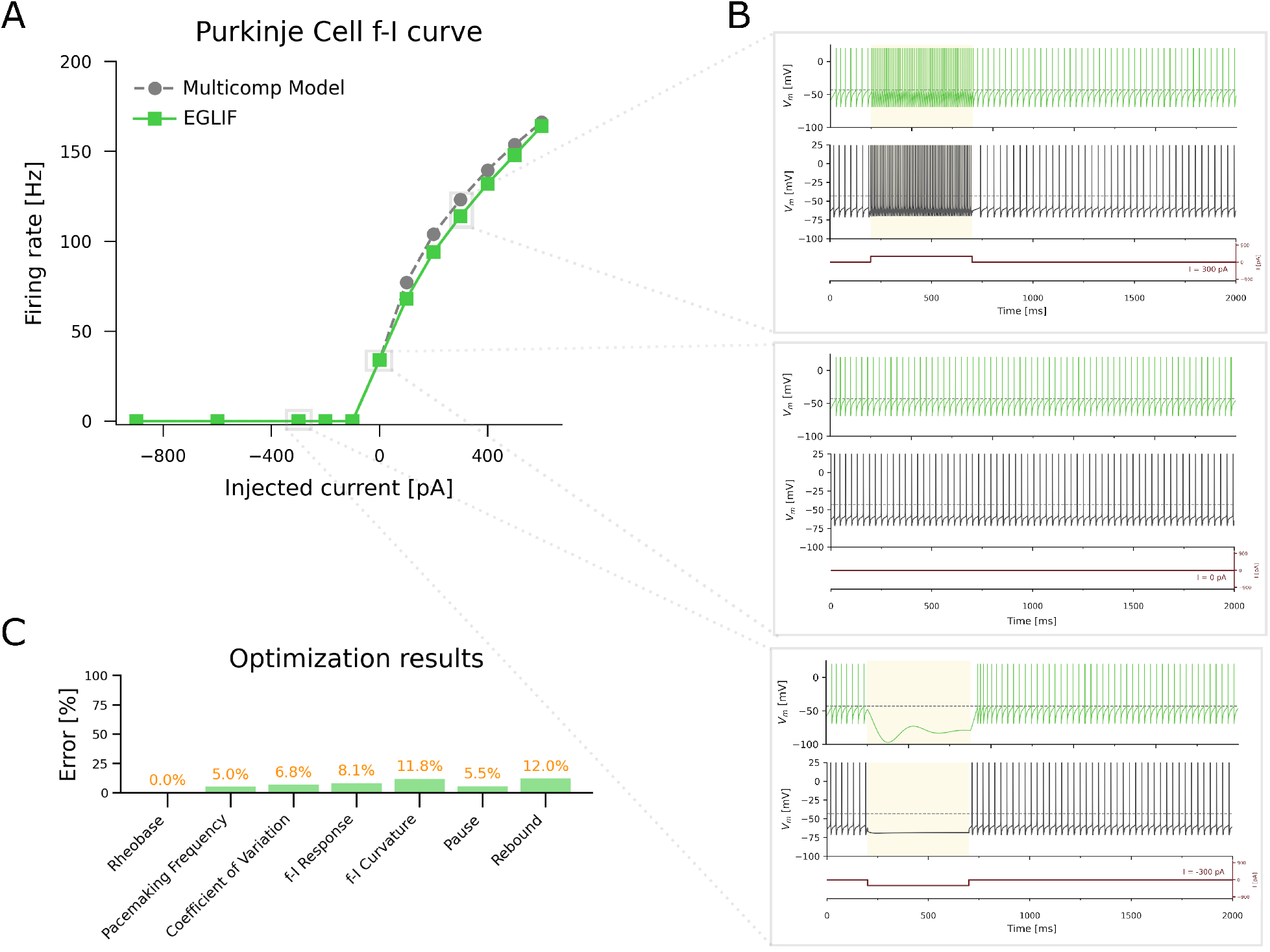
Optimization of the Purkinje cell point-neuron model. **A**. Comparison between the *f*–*I* curves of the optimized E-GLIF model and the multicompartmental reference model [10]. **B**. Representative membrane potential traces comparing the point-neuron and multicompartmental models during spontaneous activity and in response to depolarizing and hyperpolarizing current steps (*I* = 300 pA and *I* = −300 pA). **C**. Fitness errors obtained after the multi-objective optimization, reported as percentages.

The comparison was then extended to the other cerebellar cortical populations, as reported in **Figure 4**. For each cell type, current-injection ranges were chosen according to the corresponding multicompartmental reference studies [10–14], allowing the reduced models to be tested within physiologically meaningful excitability regimes. Across GrCs, GoCs, BCs, SCs, and PCs, the E-GLIF models reproduced the main population-specific input-output properties, including firing threshold and input-dependent firing modulation, supporting their use in the subsequent network-level simulations.

**Fig 4.**
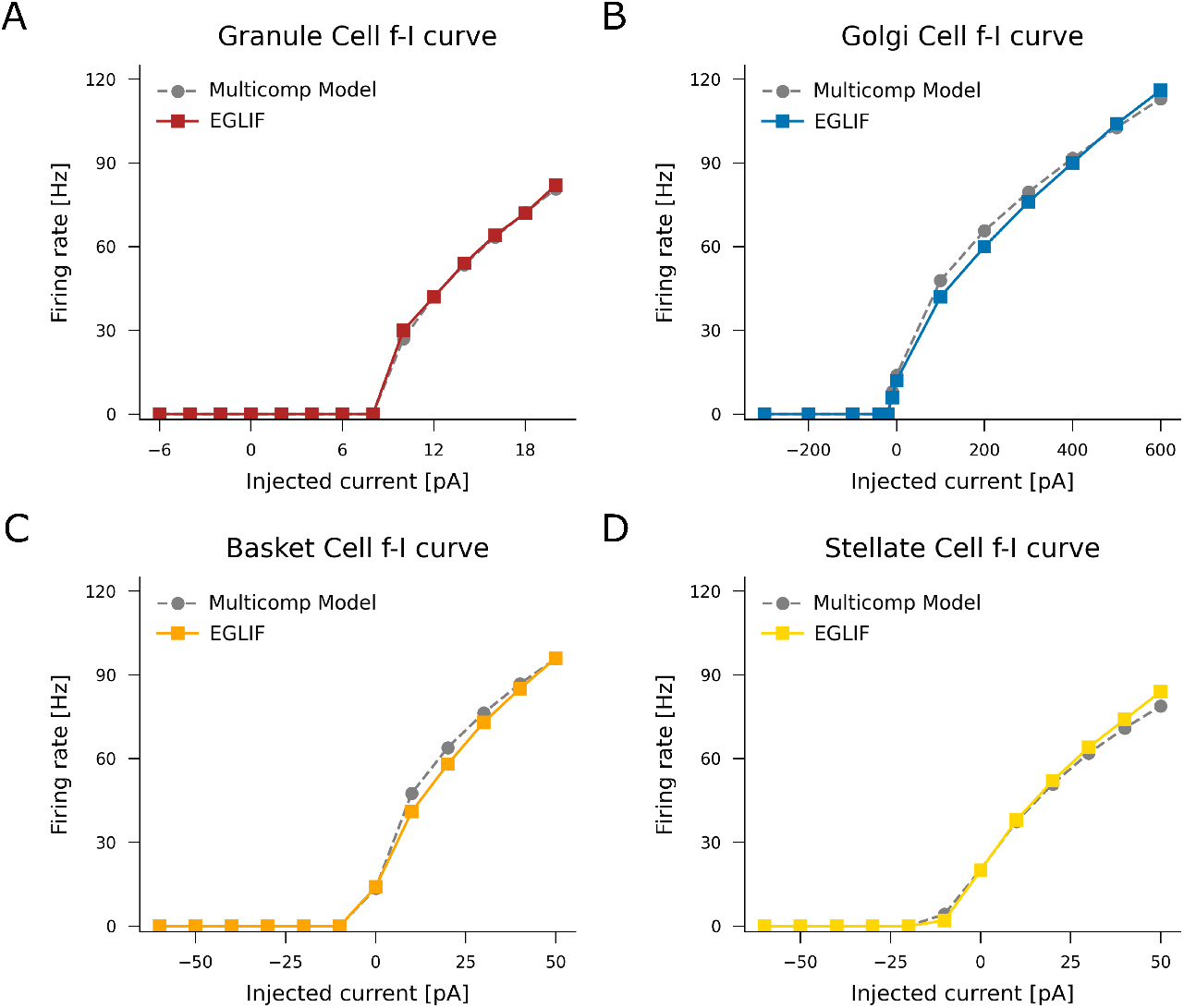
Optimized *f*–*I* curves in E-GLIF models of granule, Golgi, stellate and basket cells. **A**. Granule cell. **B**. Golgi cell. **C**. Basket cell. **D**. Stellate cell. Overall, the tuned point-neuron models accurately reproduce the *f*–*I* relationships of the multicompartment models over the explored range of injected currents, supporting their ability to capture the main input–output properties of each cell type.

Alongside the optimization of intrinsic excitability, receptor-specific synaptic transmission was described through dedicated kernels fitted to multicompartmental conductance traces.

The resulting kernels accurately reproduced the temporal structure of the reference responses across receptor types and synaptic connections. Representative examples for GrCs are shown in **Figure 5A**, where the E-GLIF kernel-based synaptic activations closely match the reference synaptic conductances simulated under voltage-clamp conditions at −40 mV, capturing both the peak amplitude and the characteristic rise and decay phases of the synaptic response.

**Fig 5.**
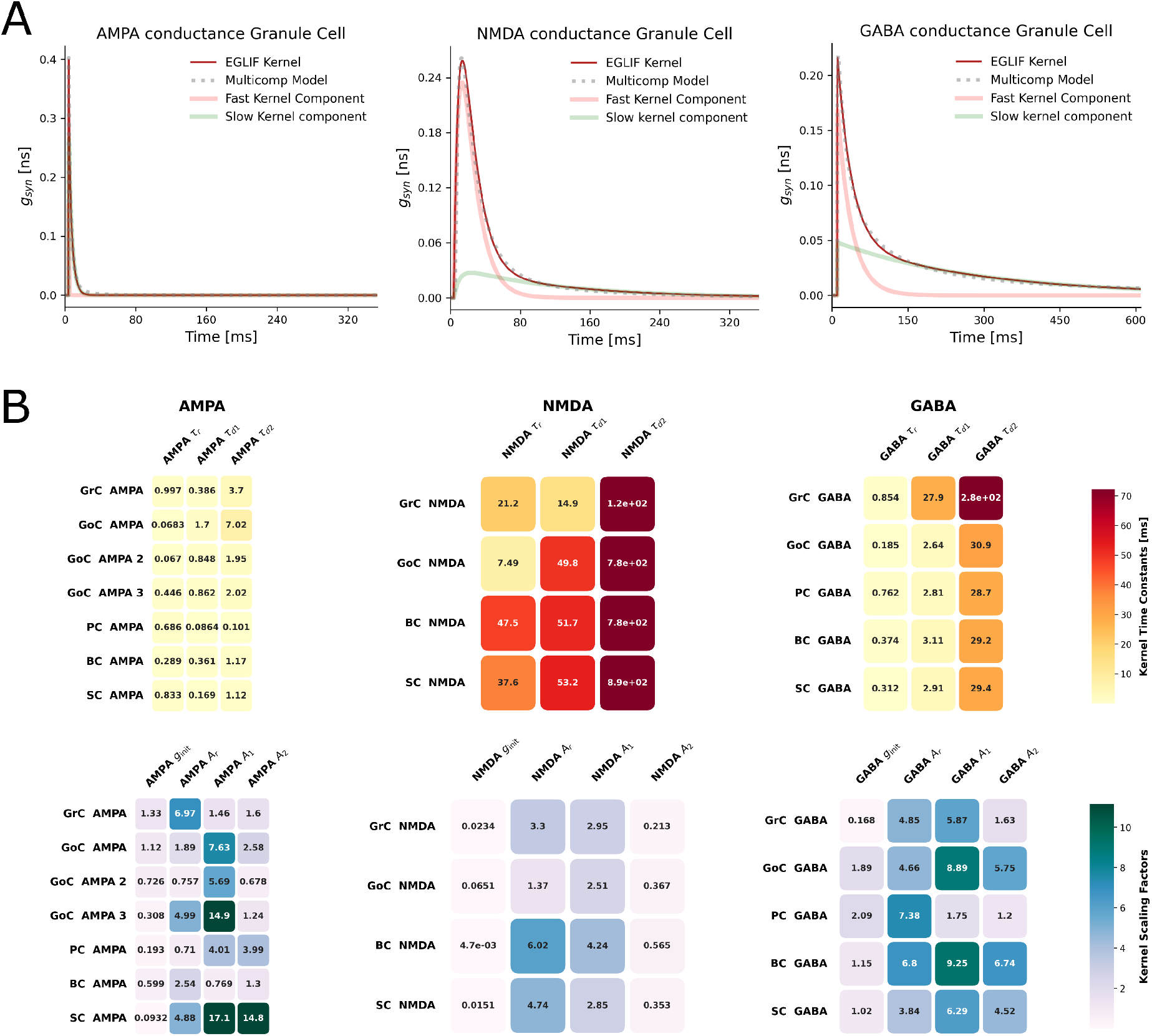
Calibration of receptor-specific synaptic kernels. **A**. Representative comparison between calibrated E-GLIF synaptic conductance kernels and reference multicompartmental conductance traces for AMPA, NMDA, and GABA receptors in granule cells. The E-GLIF kernels, decomposed into fast and slow components, reproduce the main receptor-specific features of the reference traces, including peak amplitude, rise phase, and decay kinetics. **B**. Summary of the optimized synaptic-kernel parameters across connections and postsynaptic cell types. The upper panels show the kernel time constants, while the lower panels report the corresponding scaling factors.

The full set of optimized parameters is reported in **Figure 5B** The fitted kernel time constants exhibit a clear differentiation across receptor types, reflecting their distinct synaptic dynamics. In particular, AMPA-mediated responses are characterized by almost instantaneous rising time constant (*<* 1ms) and decay constants below 10 ms, in agreement with their expected fast kinetics. In contrast, NMDA receptors display markedly slower and more heterogeneous dynamics across cell types, with decay time constants (*τ*_*d*2_) extending up to ∼900 ms. GABAergic synapses show intermediate temporal behavior between AMPA and NMDA, with slower dynamics than AMPA currents but substantially faster than NMDA-mediated ones. Among inhibitory synapses, only the granule cell GABA conductance exhibits a pronounced contribution from the second decay component, with a *τ*_*d*2_ higher than 200 ms, which significantly prolongs the inhibitory effect on GrCs.

The optimized *g*_init_ values remain below 2, whereas the amplitude parameters (*A*_*r*_, *A*_*d*1_, *A*_*d*2_) range between 0.7 and 17, with the highest values observed for AMPA-mediated synapses, consistent with the relatively large and fast conductance contribution due to AMPA receptors.

The STP parameters *U, τ*_fac_, *τ*_rec_, and *τ*_psc_ were assigned for each connection type from literature-based parameter sets [42], while only synaptic weights scaling was applied, as described in Materials and Methods. In single-cell stimulation protocols with presynaptic spike trains at different input frequencies, these connection-specific STP profiles produced postsynaptic responses dominated by either depression or facilitation. The network-level consequences of these frequency-dependent synaptic dynamics across successive mf input pulses are further investigated in the following results.

### Cerebellar SNN simulations

The final effective static and STP synaptic weights reported in Table 4 represent the outcome of the network-level calibration procedure described in Materials and Methods, aimed at reproducing the basal activity regime of the multicompartmental cerebellar circuit by De Schepper et al., 2022 [5]. The calibrated SNN was first tested under sparse mossy-fibre background drive, corresponding to the basal protocol used for network tuning. As shown in **Figure 6A**, the network reproduced population-specific spontaneous firing rates close to the reference model, with: 0.5±1 Hz for GrCs, 10±8 Hz for GoCs, 34±2 Hz for PCs, 6±9 Hz for BCs, and 9.5±13 Hz for SCs.

**Table 4.** Synaptic weights for static and short-term plasticity (STP) at the chemical synapses of the cerebellar SNN. STP weights were obtained by applying connection-specific rescaling factors, as reported in Materials and Methods.

| Connection | Receptor | Static weight | STP weight |
| --- | --- | --- | --- |
| Glomerulus → GrC | AMPA, NMDA | 1.0 | 2.4 |
| Glomerulus → GoC | AMPA, NMDA | 0.32 | 0.74 |
| GoC → GrC | GABA | 1.0 | 2.8 |
| Ascending axon (GrC) → GoC | AMPA, NMDA | 0.59 | 1.5 |
| Parallel fibers (GrC) → GoC | AMPA | 0.20 | 0.49 |
| Ascending axon (GrC) → PC | AMPA | 0.75 | 5.6 |
| Parallel fibers (GrC) → PC | AMPA | 1.24 | 9.3 |
| Parallel fibers (GrC) → SC | AMPA, NMDA | 0.51 | 3.4 |
| Parallel fibers (GrC) → BC | AMPA, NMDA | 0.29 | 1.9 |
| SC → PC | GABA | 0.14 | 0.40 |
| BC → PC | GABA | 0.47 | 1.3 |
| SC → SC | GABA | 0.035 | 0.084 |
| BC → BC | GABA | 0.016 | 0.038 |

**Fig 6.**
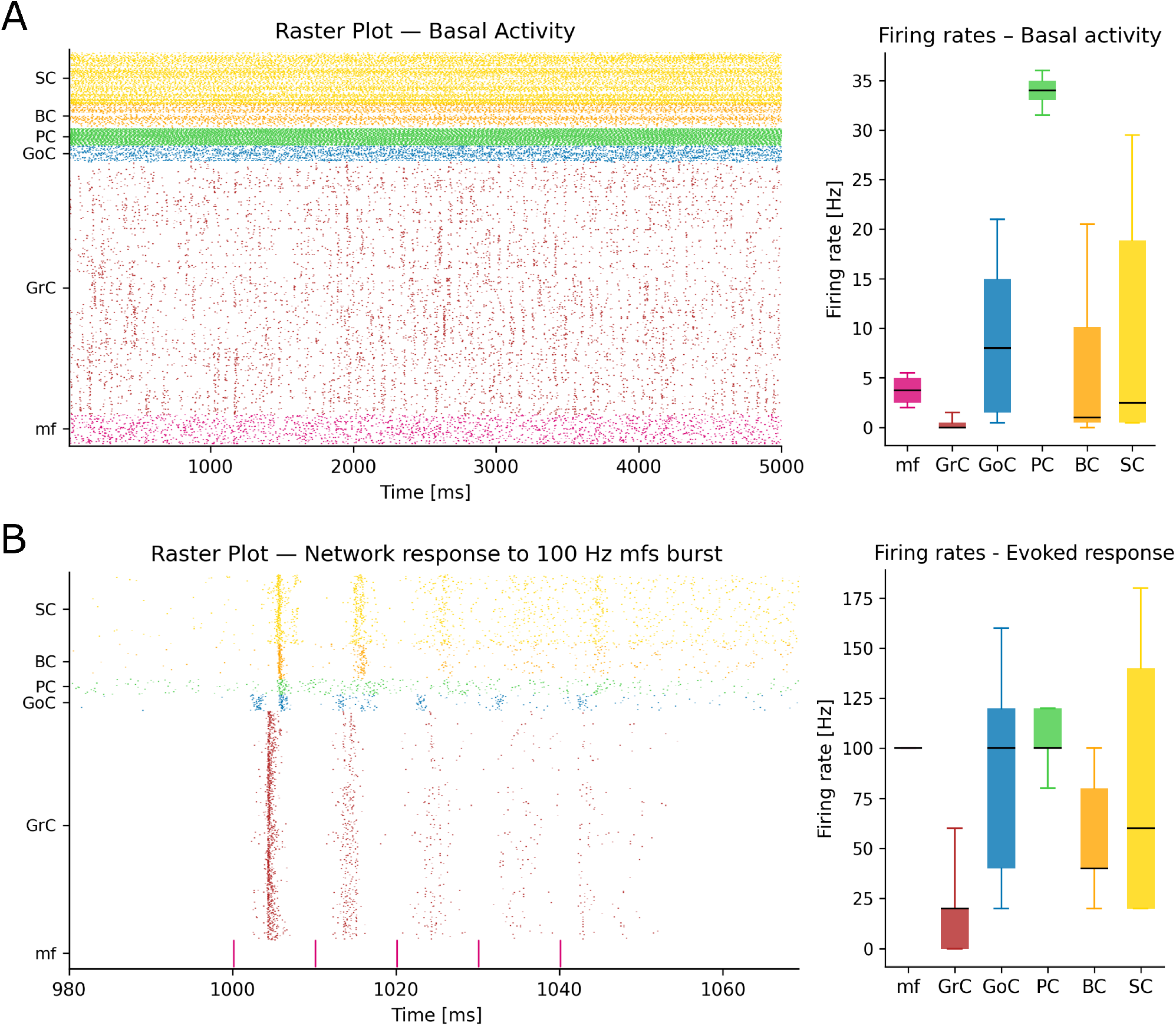
Network activity in basal conditions and during burst stimulation. **A**. Basal activity. Left: raster plot during sparse mossy-fiber background drive, modeled as a 4 Hz Poisson process on the mf population. Right: box plots of population firing rates. **B**. Evoked burst response. Left: raster plot following 5 synchronous mf pulses at 100 Hz. Right: box plots of population firing rates computed over the evoked response window [1000, 1050] ms. Mfs, GrCs, GoCs, PCs, BCs, and SCs are shown;for visual clarity, only 1000 randomly selected GrCs are displayed in the raster plots.

This basal regime provided the reference network state from which evoked responses to mossy-fiber burst stimulation were subsequently analyzed. In particular, the tuned network was first tested under evoked stimulation using a 100 Hz, 5-pulse mossy-fibre burst, as shown in **Figure 6B**. During the burst window ([1000, 1050] ms), the response propagated through the feedforward pathway and recruited local inhibition, with mean firing rates of 20±26 Hz in GrCs, 88±48 Hz in GoCs, 101±14 Hz in PCs, 58±32 Hz in BCs, and 84±64 Hz in SCs. Interestingly, activity progressively decreased across successive pulses, revealing a short-term depression pattern and indicating that STP shaped the temporal evolution of the network response under high-frequency input.

Given their central role in transforming mossy-fiber input and contributing to the simulated LFP proxy signals, GrC activity was analyzed to assess the impact of receptor-specific synaptic dynamics on granular-layer spiking. For direct comparison with the experimental conditions, responses were analyzed using a single mossy-fiber stimulation protocol without background noise, as described in Materials and Methods. This controlled setting isolated the evoked response under control and receptor-specific perturbation conditions, as shown in **Figure 7**.

**Fig 7.**
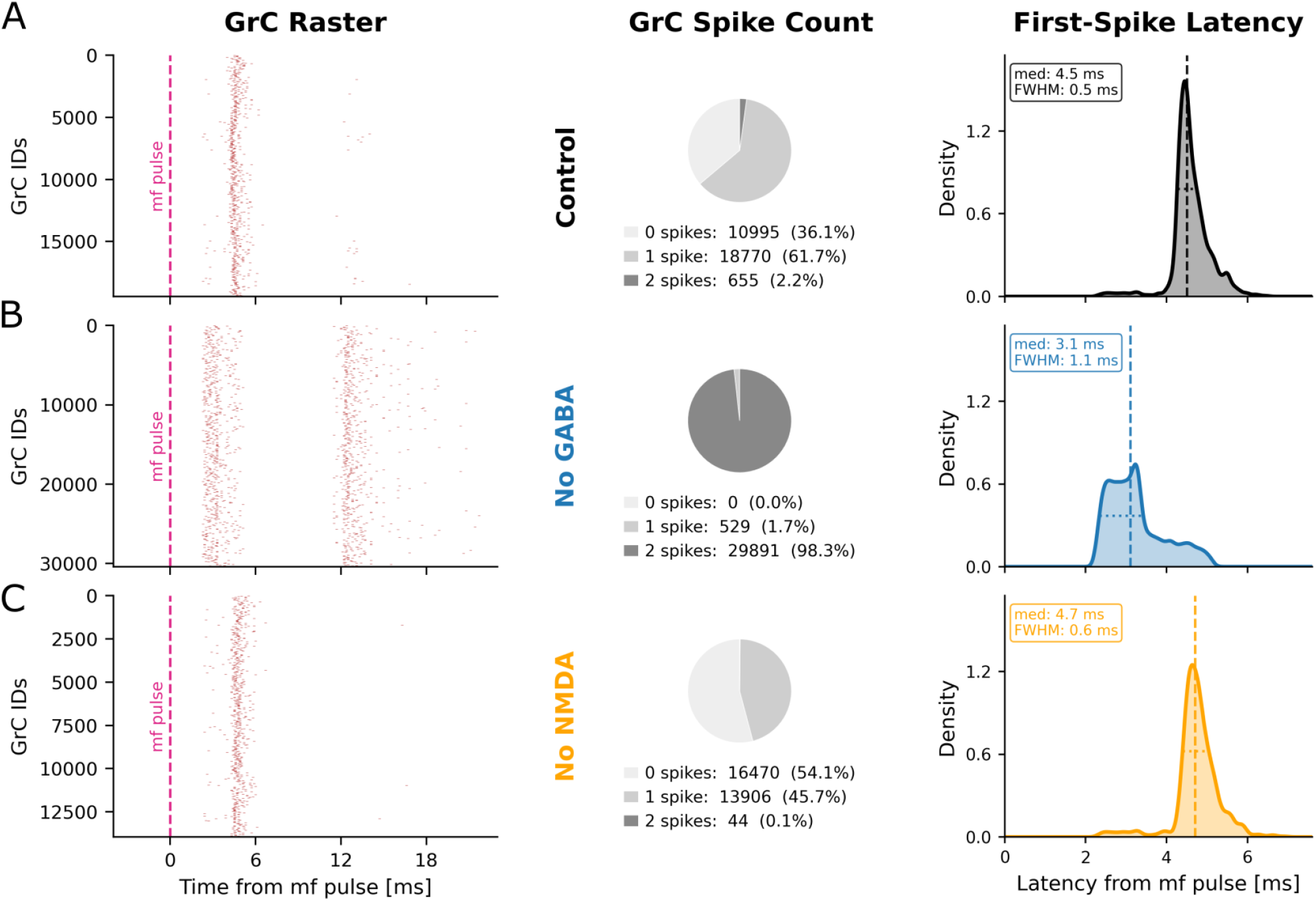
GrCs responses under control and receptor-specific blockade conditions. Spiking activity evoked in the GrC population by single mossy-fiber stimulation. For each condition, the left panel shows the raster plot, the middle panel reports the distribution of spike counts per GrC, and the right panel shows the distribution of first-spike latencies (median and Full Width at Half Maximum, FWHM). **A**. Control condition, characterized by sparse and temporally precise responses dominated by single-spike events. **B**. No GABA condition. Removal of inhibitory transmission increased GrC excitability, promoting double-spike responses and earlier first-spike latencies, with broader temporal dispersion across cells. **C**. No NMDA condition. NMDA receptor blockade reduced GrC recruitment and response persistence, increasing the fraction of silent neurons and nearly suppressing double-spike events.

We first examined the control condition (see **Figure 7A**), in which the granular layer operated in a sparse response regime: approximately 36%of GrCs remained silent, 62%generated a single spike, and only 2%produced two spikes. The first-spike latency distribution was sharply locked to the mossy-fiber input, with a median latency of 4.5 ms and a FWHM (Full Width at Half Maximum) of 0.5 ms, indicating temporally precise recruitment of the active GrC population.

GABA receptor blockade markedly increased GrC excitability (see **Figure 7B**). In this condition, almost all GrCs became active, and the response was dominated by double-spike events, rising from ∼2%in control to ∼98%. The median first-spike latency decreased to 3.1 ms, showing faster recruitment after removal of inhibition. However, the latency distribution broadened, with FWHM increasing to 1.1 ms, indicating that the enhanced excitability was accompanied by a loss of temporal precision across the population.

By contrast, NMDA receptor blockade shifted the network toward a less active regime. The fraction of silent GrCs increased to ∼54%, while double-spike responses were almost completely suppressed. The median first-spike latency remained close to control values, 4.7 ms, with a narrow FWHM of 0.6 ms. Overall, these results indicate that granular-layer output is shaped by the balance between NMDA-dependent recruitment of GrCs and GABA-mediated control of response gain, timing, and burst-like activity.

After characterizing receptor-dependent changes in GrC spiking activity, we examined whether these effects were reflected in mesoscopic LFP-like signals. The LFP proxies were used as population-level readouts of granular-layer synaptic activity, enabling comparison between simulated circuit dynamics and HD-MEA recordings.

The analysis pipeline used for the internal HD-MEA LFP datasets is illustrated in **Figure 8A** through a representative experiment. In this slice, 58 valid channels were retained after reliable detection of both N2A and N2B peaks. The resulting peak features showed the expected organization of the granular-layer LFP response, with N2A characterized by larger amplitude and shorter latency than N2B. These experimentally extracted descriptors were then used as benchmarks for the comparison with the simulated LFP proxy signals. To generate the corresponding model-based readout, a single mossy-fiber bundle pulse was delivered to the cerebellar SNN. Fifteen equispaced virtual channels were placed over the simulated granular layer, each sampling a local group of GrCs (16±4 neurons per channel). Given the limited spatial extent of the simulated network, these channels were interpreted as local sampling regions surrounding the stimulation site, rather than as a full reconstruction of the broader experimental recording field.

**Fig 8.**
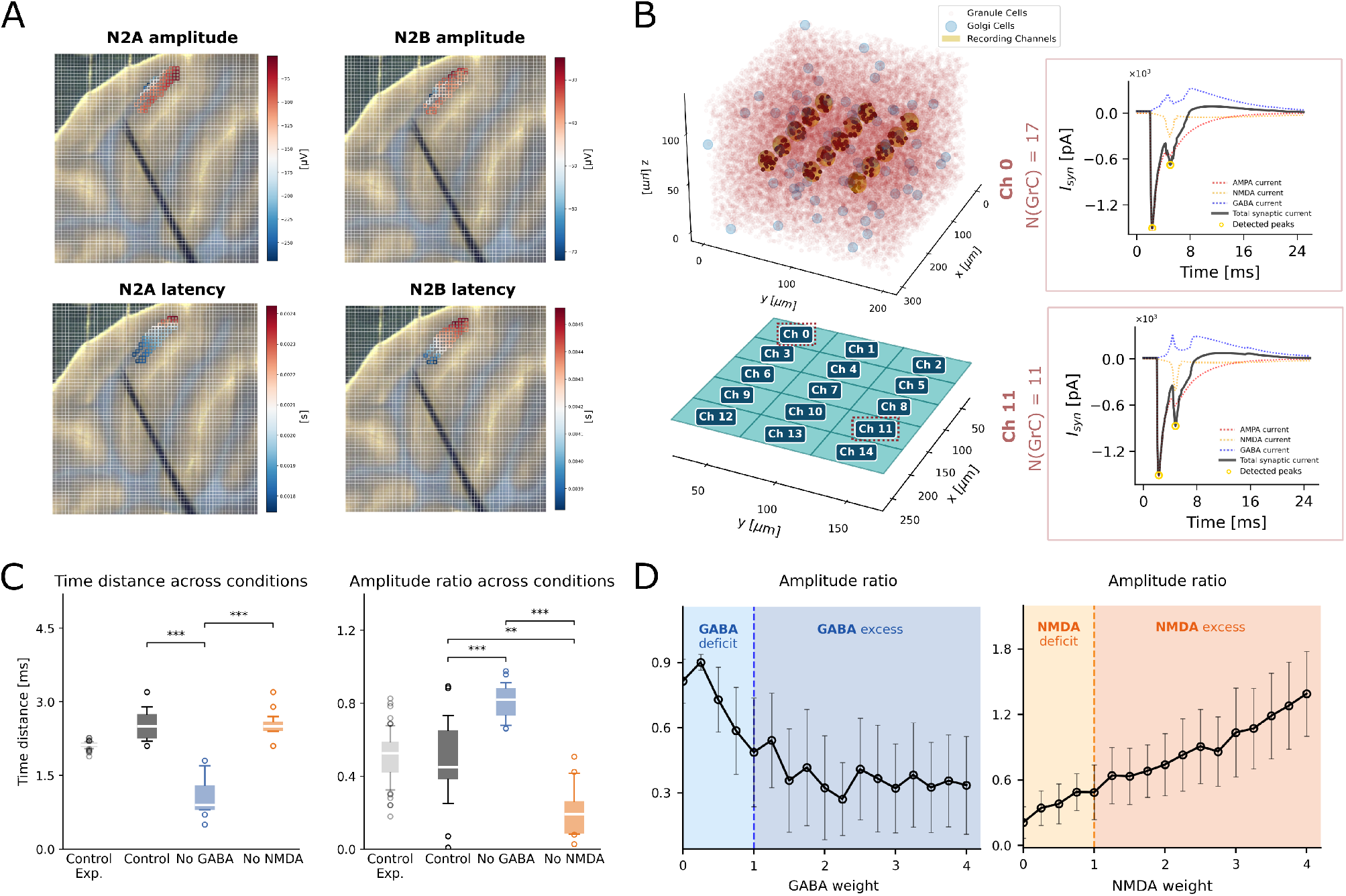
Experimental and simulated granular-layer LFP features and receptor-specific contributions. **A**. Representative spatial maps of N2A and N2B amplitudes and latencies extracted from HD-MEA recordings after mossy-fiber stimulation, overlaid on the recorded slice and stimulation electrode position, showing the localized activation pattern of the granular layer. **B**. Simulated cerebellar SNN with virtual recording channels positioned within the modeled granular-layer volume. Representative LFP proxy traces were computed from the summed synaptic currents generated by GrCs within each sampling volume and decomposed into AMPA, NMDA, and GABA components. **C**. Quantitative comparison of LFP peak features across experimental control, simulated control, and receptor-specific blockade conditions. Left: temporal distance between N2A and N2B. Right: N2B/N2A amplitude ratio. **D**. Sensitivity analysis of the N2B/N2A amplitude ratio to receptor-specific synaptic scaling. NMDA scaling was applied to Glom→GrC excitation, whereas GABA scaling was applied to GoC→GrC inhibition. Error bars indicate variability across virtual channels.

Within each virtual channel, the summed synaptic activity of the sampled GrCs produced LFP proxy traces with identifiable N2A and N2B components, enabling the same peak-based analysis applied to the experimental recordings. Representative examples are shown in **Figure 8B**. In these traces, the synaptic-current decomposition highlights the prominent contribution of NMDA-mediated currents to the delayed N2B component, whereas channel-dependent variability reflects local differences in microcircuit organization, specifically the heterogeneous recruitment of GoC-mediated inhibition.

The quantitative comparison between experimental and simulated responses is reported in **Figure 8C**. Under control conditions, the simulated LFP proxies fell within the experimental range for both the N2B/N2A amplitude ratio and the inter-peak temporal distance, as further summarized in Table 5. This agreement indicates that the calibrated SNN reproduces not only the spike-based output of the granular layer, but also the synaptic population dynamics underlying the experimentally observed LFP components.

**Table 5.**
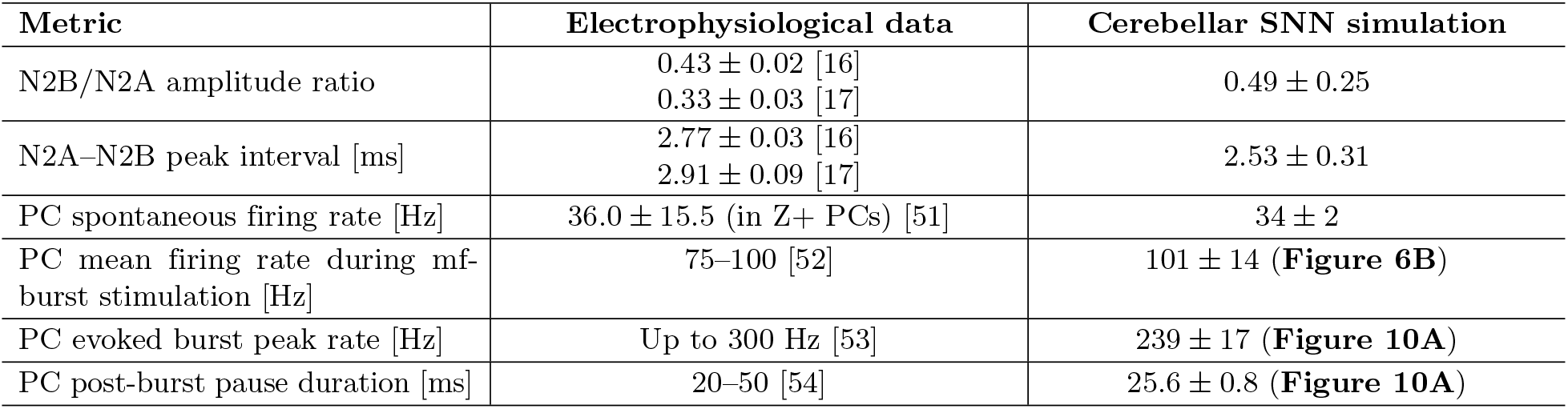
Comparison between electrophysiological data from experimental recordings or literature references and cerebellar SNN simulations for granular layer LFP-derived metrics and Purkinje cell activity features. For Purkinje cells, spontaneous activity was compared with simple-spike firing in zebrin-positive cells. Evoked activity was assessed using the mean firing rate during the 100 Hz 5-pulse mf protocol, whereas the PC evoked peak and post-burst pause duration were derived from the PC response to a single mf pulse.

The same receptor-specific perturbations analyzed at the spiking level in Figure 7 were then evaluated at the level of the simulated LFP proxies. In the simulations, GABA receptor blockade increased the N2B/N2A amplitude ratio (0.8±0.1), mainly by enhancing the second peak, and markedly shortened the N2A–N2B inter-peak interval to 1.1±0.4 ms. This is consistent with the more excitable and burst-prone GrC regime observed after removal of inhibition. By contrast, NMDA receptor blockade preserved an inter-peak distance close to control values (2.5±0.3 ms), but strongly reduced the relative amplitude of N2B, yielding an amplitude ratio of 0.2±0.1. This agrees with the reduced GrC recruitment and the suppression of double-spike responses observed when NMDA-mediated transmission was removed.

Specifically, pairwise two-sample Kolmogorov–Smirnov tests revealed that GABA blockade significantly shifted the N2B–N2A time distance relative to Control (*KS* = 1.000, *p* = 3.87×10^−8^) and NMDA blockade (*KS* = 1.000, *p* = 1.07×10^−7^), while Control and NMDA blockade did not differ (*p* = 0.696). The amplitude ratio differed across all pairs: Control vs. GABA blockade (*KS* = 0.733, *p* = 7.07×10^−4^), Control vs. NMDA blockade (*KS* = 0.590, *p* = 0.008), and GABA vs. NMDA blockade (*KS* = 1.000, *p* = 1.60×10^−7^).

To further assess the dependence of the LFP proxies on receptor-specific transmission, NMDA and GABA synaptic strengths were systematically varied by applying a multiplicative scaling factor in the range [0, 4], with 1 corresponding to the control condition (see **Figure 8D**). The scaling was respectively applied to Glom→GrC for NMDA-mediated excitation and to GoC→GrC for GABA-mediated inhibition. This analysis showed that the N2B/N2A amplitude ratio increased with NMDA strength, reaching values above 1 under strong NMDA amplification, indicating that the second peak can become dominant when slow excitatory integration is enhanced. Conversely, increasing GABAergic weight progressively suppressed the second peak, with saturation for inhibitory strengths above approximately twice the control value.

Overall, these results indicate that the simulated N2A–N2B structure reflects the dynamical state of the granular-layer circuit. NMDA-mediated transmission supports the delayed N2B component, whereas GABAergic inhibition constrains both response amplitude and timing. Thus, the LFP proxy links receptor-dependent synaptic mechanisms to mesoscopic population signals comparable to experimental recordings.

After the initial assessment based on the single mossy-fiber stimulation protocol, we investigated how short-term plasticity shaped LFP proxy dynamics during repeated stimulation. Simulated LFP proxies were recorded during 5-pulse mossy-fiber bursts delivered at different frequencies, as described in Materials and Methods. In **Figure 9**, we report the subset of frequencies {6, 20, 50, 100} Hz, selected to match the experimental LFP protocols used as reference [16, 17].

**Fig 9.**
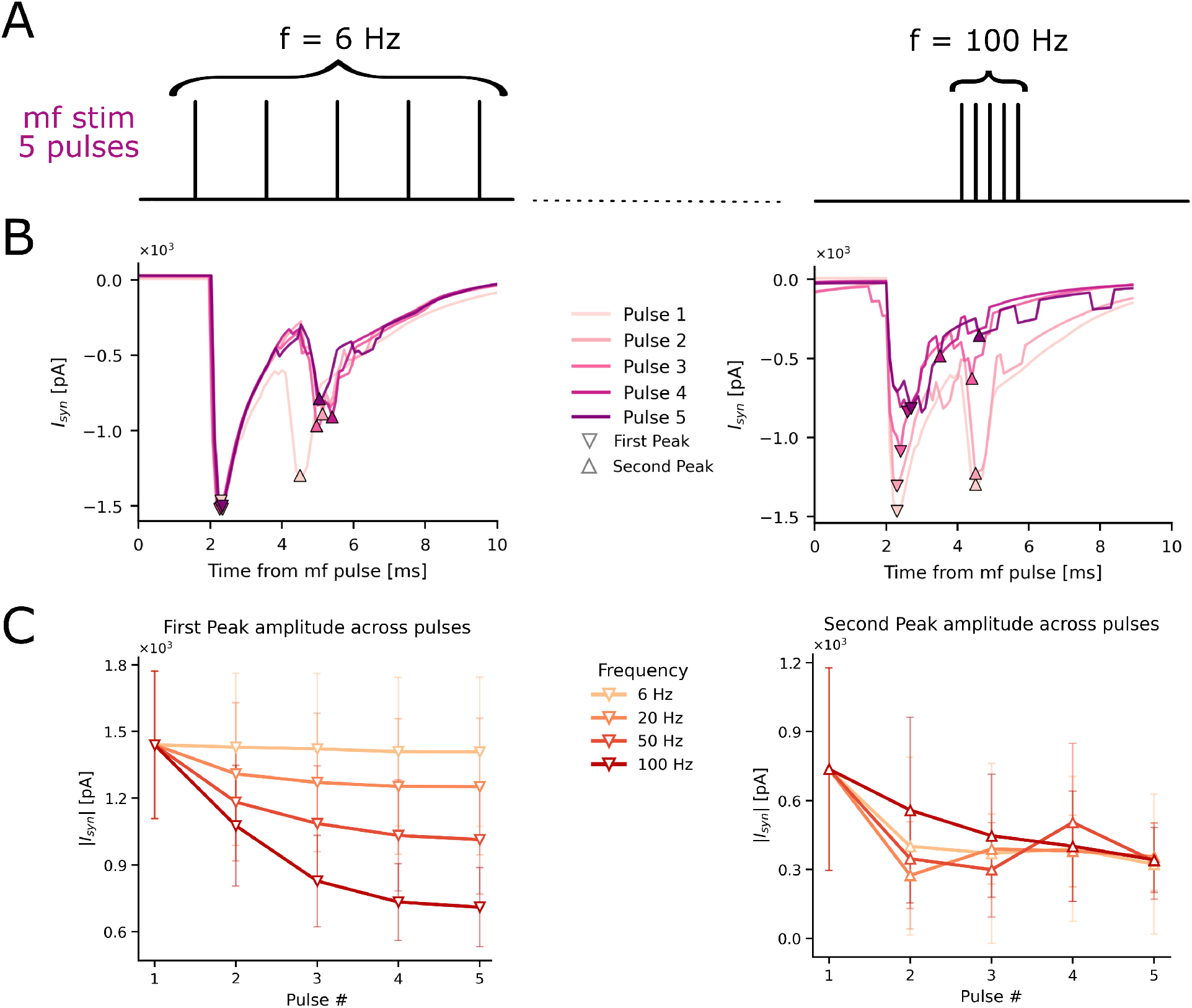
Short-term plasticity effects on LFP proxies during mf stimulation protocols at different frequencies. **A**. Schematic representation of the stimulation protocol, consisting of 5-pulse trains delivered at different frequencies, with examples shown for 6 Hz and 100 Hz. **B**. Representative LFP proxy responses across the 5 pulses at low (6 Hz) and high (100 Hz) stimulation frequencies, illustrating the frequency-dependent modulation of the first and second peaks. **C**. Quantitative analysis of peak amplitudes across pulses for the subset of stimulation frequencies considered in the LFP comparison (6, 20, 50, and 100 Hz). Left: first peak amplitude as a function of pulse number. Right: second peak amplitude as a function of pulse number. The first peak shows progressively stronger depression at increasing stimulation frequency, whereas the second peak displays a more complex pulse-dependent modulation.

The LFP proxy signals exhibited frequency-dependent modulation of peak amplitudes across pulses. At low frequency, the first peak of the synaptic response remained largely stable across pulses, whereas the second peak showed a mild reduction, mainly between the first and subsequent pulses. With increasing input frequency, the first peak exhibited more pronounced short-term depression, with an approximately exponential decay across pulses, consistent with the adopted Tsodyks–Markram parametrization of granular-layer synapses. The second peak also showed pulse-dependent modulation, although without a monotonic frequency-dependent trend. This behavior is expected, since the delayed component does not reflect STP alone, but rather the combined effect of short-term synaptic dynamics, local inhibition, and GoC-mediated feedback within the granular-layer circuit.

Finally, PC spiking activity was analyzed as the final readout of cerebellar cortical dynamics. Under baseline conditions, the simulated PC population fired spontaneously at 34±2 Hz, also consistent with in vivo spontaneous firing rates reported for zebrin-positive Purkinje cells (Z+PCs) (see Table 5).

We then examined the PC population response to a single mf pulse under control conditions and after removal of MLI-mediated inhibition, obtained by deactivating the SC→PC and BC→PC synapses (see **Figure 10A**). In control conditions, the mf input evoked a burst-pause response, with a rapid firing-rate peak of 239±17 Hz followed by a pronounced suppression phase. The pause lasted 25.6±0.8 ms, during which PC firing dropped close to silent levels before gradually returning to baseline through damped oscillatory activity. These response features are consistent with experimental findings on Purkinje cell activity recorded after mf stimulation using comparable protocols, as summarized in Table 5.

**Fig 10.**
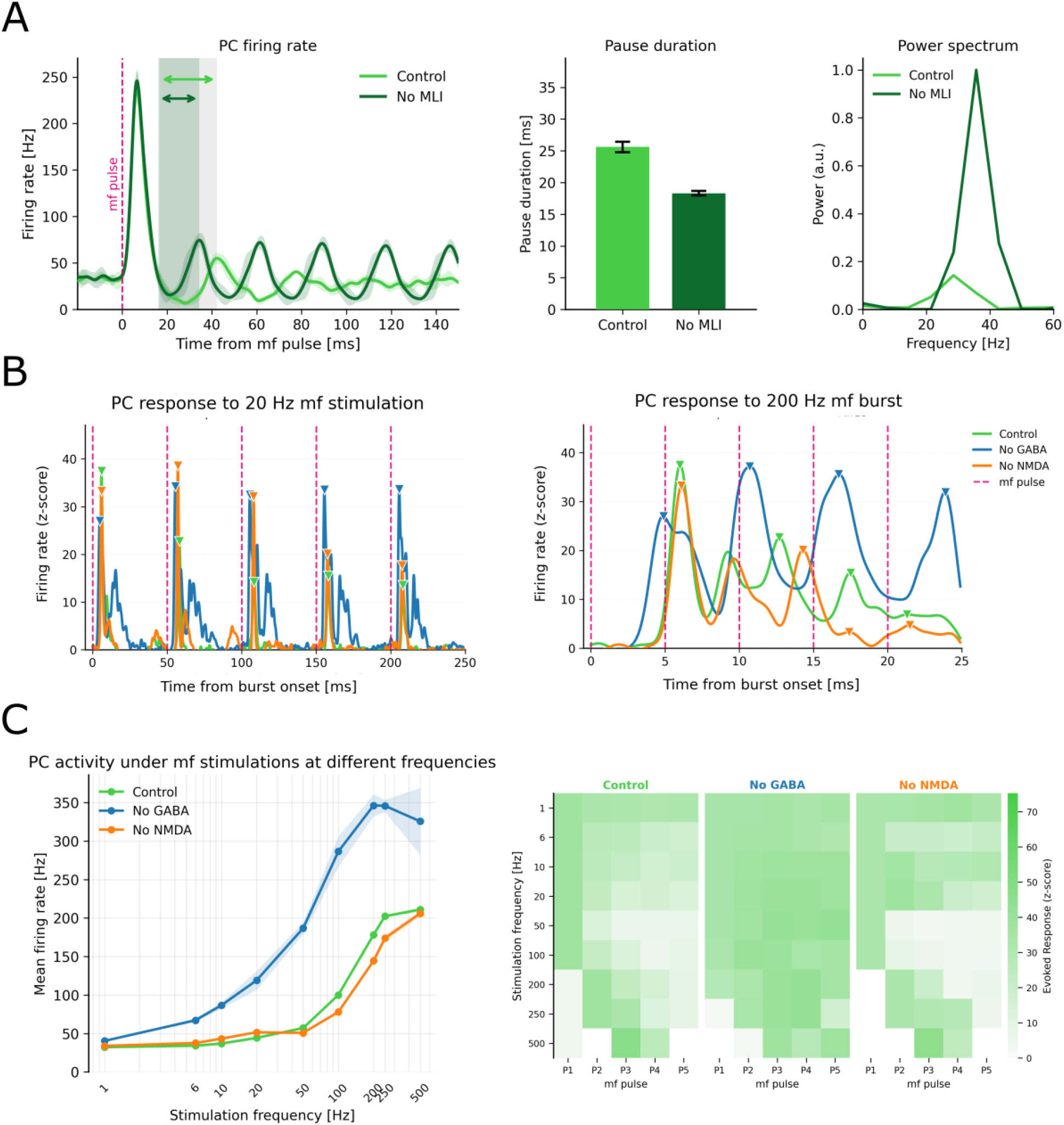
Simulated Purkinje cell population responses to mf stimulation protocols. **A**. PC population response to a single mf pulse under control conditions and after removal of MLI-mediated inhibition. Left: instantaneous PC population firing rate, estimated with a Gaussian kernel (*σ* = 2 ms), with shaded regions indicating the post-burst pause intervals. Middle: post-burst pause duration, measured from the first drop below baseline after the evoked peak to the subsequent local maximum. Right: corresponding post-stimulus power spectra. **B**. Representative PC population responses to 5-pulse mf stimulation at 20 Hz and 200 Hz under Control, No GABA, and No NMDA conditions. **C**. Frequency-dependent PC responses during repeated mf stimulation. Left: mean PC firing rate computed over the full stimulation window for each input frequency. Right: pulse-resolved heatmaps reporting the evoked response associated with each mf pulse across frequencies and receptor conditions. For panels B–C, firing rates were estimated with a Gaussian kernel (*σ* = 0.5 ms), kept constant across frequencies to allow pulse-resolved comparison.

Removing MLI input preserved the initial evoked excitation but markedly reduced the subsequent suppression phase, shortening the pause to 18.3±0.4 ms. After the stimulus, PC activity rapidly settled into a synchronized oscillatory pattern, suggesting that the mf pulse transiently reset the phase of the population pacemaking rhythm. Consistently, the post-stimulus power spectrum in the absence of MLI inhibition was dominated by a component close to the intrinsic PC autorhythmic frequency, with a peak at 36 Hz. By contrast, under control conditions, MLI-mediated inhibition limited the emergence of this synchronized post-stimulus rhythm, reducing the spectral peak and slightly shifting the dominant frequency toward lower values.

These results suggest that MLI-mediated GABAergic inhibition may limit the phase-reset-like synchronization of PC activity after mf stimulation, reducing the persistence of post-stimulus oscillations in the population firing rate.

We extended the analysis of PC population activity to repetitive mf stimulation protocols, assessing how the final cerebellar cortical output evolved across consecutive inputs in the presence of receptor-specific synaptic dynamics, STP, and circuit-level inhibition. Representative responses to 5-pulse mf stimulation delivered at 20 Hz and 200 Hz are shown in **Figure 10B**.

At 20 Hz, the control response showed a progressive reduction across the 5 mf stimuli, reflecting the combined effect of short-term synaptic dynamics, receptor-specific kinetics, and inhibitory feedback within the network. Under NMDA blockade, this pulse-by-pulse depression was less pronounced, suggesting that NMDA removal affects both GrC activation and the recruitment of inhibitory pathways, thereby altering the excitation/inhibition balance onto the PC population and reshaping its dynamical response. While, GABA blockade increased PC excitability and prolonged the post-pulse response, reflecting the loss of inhibitory control that normally sharpens GrC-driven excitation and subsequently PC firing.

At 200 Hz, the short inter-pulse interval prevented PC activity from returning to baseline, leading to temporal summation of the evoked responses. In this regime, NMDA blockade altered the temporal buildup of activity, whereas GABA blockade strongly increased and prolonged the population response.

These frequency-dependent responses are further summarized in **Figure 10C**. The mean PC firing rate was computed over the full stimulation window associated with each input frequency;for example, the 100 Hz mf burst corresponded to a 50 ms window and yielded a mean PC firing rate of 101±14 Hz, similarly to the representative network response of Figure 6B. Across frequencies, the mean PC output increased nonlinearly with stimulation frequency. In control conditions, PC activity remained relatively close to baseline at low frequencies and then rose sharply above 50–100 Hz, reaching a plateau at the highest frequencies. Blocking GABAergic inhibition strongly increased the gain of this frequency-dependent response, producing higher mean firing rates across the whole frequency range and an earlier saturation at high-frequency stimulation. Conversely, NMDA blockade reduced the high-frequency buildup of PC activity, delaying the increase in mean firing rate and yielding lower responses than control at intermediate-to-high frequencies. The pulse-resolved heatmaps provide a compact summary of the response dynamics across the 5 mf stimuli, showing how the trends observed in the representative examples generalize across the full stimulation set.

## Discussion

The model developed in this work is proposed as a canonical point-neuron SNN of the cerebellar cortex, embedding fundamental neuronal and synaptic mechanisms required to simulate complex spatiotemporal circuit dynamics. It therefore bridges the gap between multicompartmental models, which offer high mechanistic precision but are computationally expensive and poorly scalable for massive simulations, and abstract point-neuron models, which are efficient but often lack key biophysical and synaptic mechanisms.

Classical theories of cerebellar computation, starting from Marr and Albus [55, 56], provided a foundational description of information transfer and learning across cerebellar layers, later extended within adaptive-filter frameworks [57]. However, these theoretical formulations necessarily abstracted from the nonlinear excitable properties of neurons and from receptor-specific synaptic mechanisms. Subsequent computational models progressively introduced single-neuron resolution, with different compromises between biological detail and scalability.

Cerebellar circuit models now span a broad range of biological detail, from multicompartmental granular-layer models with biophysical synapses [58, 59], to simplified LIF-based descriptions of granular-layer computation [60, 61], and scalable SNNs of the cerebellar cortex or selected subcircuits [62, 63]. These approaches demonstrated the computational value of cerebellar network models, but generally relied on simplified neuronal or synaptic formulations and did not include the combination of receptor-specific dynamics, voltage-dependent NMDA transmission, and short-term plasticity mechanisms considered here.

At the full cerebellar cortical scale, detailed multicompartmental models have provided a reference anatomical and biophysical reconstruction of the microcircuit [5], but remain computationally demanding for parameter sweeps and closed-loop applications. Some attempts to recover the neuronal input-output function nonlinearity into cerebellar point-neuron models have appeared [21, 27], but still relied on simplified static conductance-based synapses, excluding voltage-dependent NMDA transmission and connection-specific short-term plasticity.

The present model addresses these limitations by leveraging the E-GLIF formalism to build point-neuron models that embed key properties of intrinsic excitability and synaptic transmission. Taking detailed multi-compartmental neurons and the reconstructed cerebellar microcircuit as reference models [5], we designed an automatic calibration pipeline to generate a point-neuron network that preserves the main electrophysiological and synaptic features of the different cerebellar neuronal populations.

Specifically, biological realism was introduced through four main constructive elements: (1) cell-specific nonlinear input-output relationships and firing patterns, (2) receptor-specific synaptic conductance kernels including voltage-dependent NMDA gating, (3) connection-specific short-term plasticity, and (4) network-level effective synaptic weights.

At the single-cell level, the automatic parameter optimization pipeline generated cell-specific E-GLIF models that closely matched the main firing properties of the corresponding multicompartmental neurons. Despite their reduced point-neuron formulation, the optimized models captured complex nonlinear excitability features with limited errors across the electrophysiological targets considered in the optimization.

At the synaptic level, receptor-specific kernels enabled the model to capture synaptic conductance kinetics beyond the standard generalized alpha- or exponential-function descriptions commonly adopted in SNNs. The optimized kernels reproduced the main rise and decay dynamics of the reference conductance traces, while preserving distinct temporal and amplitude profiles for AMPA-, NMDA-, and GABA-mediated transmission [58, 64]. Moreover, the inclusion of voltage-dependent NMDA receptor gating introduced an essential nonlinear synaptic mechanism known to shape cerebellar circuit dynamics [18, 65].

The introduction of synapse-specific short-term plasticity [42, 64, 66] further enabled the model to capture frequency-dependent modulation of synaptic efficacy across cerebellar cortical pathways. Through connection-specific facilitation and depression profiles, STP shaped the temporal evolution of network responses, affecting both granular-layer dynamics and Purkinje cell output.

Moreover, when moving from multicompartmental neurons to point-neuron representations, nonlinear dendritic processing is necessarily lost [67]. In the present model, we attempted to partially compensate for this loss by optimizing synaptic weights to match the spontaneous background activity of the reference network. This can be interpreted as a first attempt to summarize, in a single connection-level parameter, the complex communication between synapses and soma, which mainly arises from dendritic attenuation, active dendritic mechanisms, and voltage-dependent conductances. Similar effective corrections have been proposed to compensate for dendritic filtering in automated point-neuron simplification workflows [47].

The resulting cerebellar SNN reproduced key properties of recordings obtained from acute cerebellar slices. First, the network model captured the main features of granular-layer evoked responses and provided a mechanistic interpretation of the relative contributions of different receptor systems to the LFP waveform [68, 69]. Although simplified, the LFP proxy reproduced the dominant experimental features, including the characteristic N2A–N2B structure observed in slice recordings. This allowed the simulated extracellular signal to be linked to specific synaptic current components and to the underlying GrC spike dynamics, providing a mechanistic bridge between receptor-dependent synaptic activity and mesoscopic LFP-like responses.

Secondly, the network model captured key features of Purkinje cell activity, including spontaneous autorhythmic firing and the burst-pause response evoked by mossy-fiber stimulation, in line with reported in vivo PC firing properties [52]. In the model, the near-silent post-burst phase emerged from the combined action of intrinsic PC excitability and feedforward inhibition mediated by molecular layer interneurons recruited by GrC activity, consistently with the role of MLI–PC interactions in shaping PC firing regularity and spike timing [63]. This coordinated pause may be relevant for cerebellar output, since synchronized pauses across converging PCs can transiently reduce inhibition onto deep cerebellar nuclei and promote time-locked nuclear spiking [54, 70]. Moreover, the comparison between control and No-MLI conditions suggests that MLI-mediated inhibition limits stimulus-induced synchronization within the PC population, promoting temporal diversity in PC output and preventing the emergence of an overly synchronized post-stimulus rhythm.

In this context, the framework can be employed to explore how receptor-mediated synaptic components influence cerebellar circuit dynamics. Selective modulation of receptor-mediated transmission allowed us to disentangle the contribution of individual synaptic currents and quantify their impact across cerebellar layers. In particular, GABA receptor switch-off produced marked alterations in both the N2A–N2B structure of granular-layer LFPs and in the dynamics of the Purkinje cell population response [69]. NMDA receptor switch-off also produced significant effects, mainly at the granular-layer LFP level, with changes opposite to those observed after GABA receptor switch-off. Together, these perturbations revealed complementary receptor-specific roles, with NMDA-mediated transmission supporting the delayed N2B component and GABAergic inhibition constraining response amplitude and timing. The gradual modulation of NMDA and GABA strengths further confirmed this balance, showing that the N2B/N2A amplitude ratio increased with NMDA-mediated excitation and decreased with stronger GABAergic inhibition. This highlights the sensitivity of granular-layer processing to receptor-specific synaptic perturbations.

Additionally, the inclusion of short-term plasticity enabled frequency-dependent modulation of the network responses. Under repeated mossy-fiber stimulation, LFP proxy components and PC firing evolved across pulses in a manner consistent with the expected facilitation and depression regimes of cerebellar synapses. These dynamics were further shaped by NMDA- and GABA-mediated transmission, reflecting the interaction between short-term synaptic plasticity, receptor-specific kinetics, and circuit-level inhibition.

Overall, the incorporation of a rich repertoire of neuronal and synaptic mechanisms allowed complex nonlinear network responses to emerge under frequency-modulated inputs, reproducing key features reported in experimental recordings of cerebellar cortical activity, including voltage-sensitive dye imaging and high-density multielectrode array recordings [16–18].

Beyond validation against physiological recordings, the same mechanistic structure makes the model suitable for exploring disease-relevant alterations of cerebellar processing. In particular, receptor-specific mechanisms represent a natural target for pathological and pharmacological simulations, since changes in synaptic conductance, kinetics, or excitation–inhibition balance can be introduced selectively and followed across multiple levels of organization.

NMDA receptor dysfunction provides a relevant example. NMDA receptor hypofunction has been implicated in schizophrenia-related synaptic deficits and altered excitation–inhibition balance [71, 72], whereas increased NMDA conductance has been reported in the IB2-KO mouse model of Phelan–McDermid syndrome, a neurodevelopmental condition associated with autism-spectrum phenotypes [73].

The proposed framework offers a controlled computational setting in which such receptor-specific alterations can be imposed at the synaptic level and propagated to network readouts, including spike timing, LFP-like signals, and Purkinje cell output. In this way, the model could be used not only to reproduce experimental perturbations, such as receptor blockade or modulation, but also to generate predictions on how pathological synaptic changes reshape cerebellar cortical dynamics and how pharmacological interventions may restore physiological activity patterns [72].

Another natural extension of the framework would be to incorporate longer-term plasticity mechanisms, such as spike-timing-dependent plasticity (STDP) or Bienenstock–Cooper–Munro (BCM) learning [6, 74, 75]. This would allow the model to investigate how receptor-specific modulation produces persistent changes in cerebellar circuit dynamics, with potential relevance for closed-loop applications and behavioural studies.

In conclusion, the proposed approach provides an effective compromise between biological realism and computational efficiency. Compared with a multicompartmental network of similar size, the point-neuron implementation substantially reduced the computational cost while preserving key cell-specific, synaptic, and network-level properties. Consistently with recent efforts to develop open modeling frameworks and structured pipelines for biologically grounded SNNs in other brain circuits [44, 76], the model and codebase presented here provide a reusable platform for studying cerebellar microcircuit dynamics, interpreting experimental recordings, and deriving reduced yet physiologically informed descriptions that can be embedded into large-scale whole-brain simulators [77] or real-time control architectures [7].

## Code Availability

All code developed for this study is publicly available in the following GitHub repository: https://github.com/dbbs-lab/cerebellar-models/tree/feature/AMPA_NMDA_GABA_eglif.

## Supporting information

**S1 Table.**
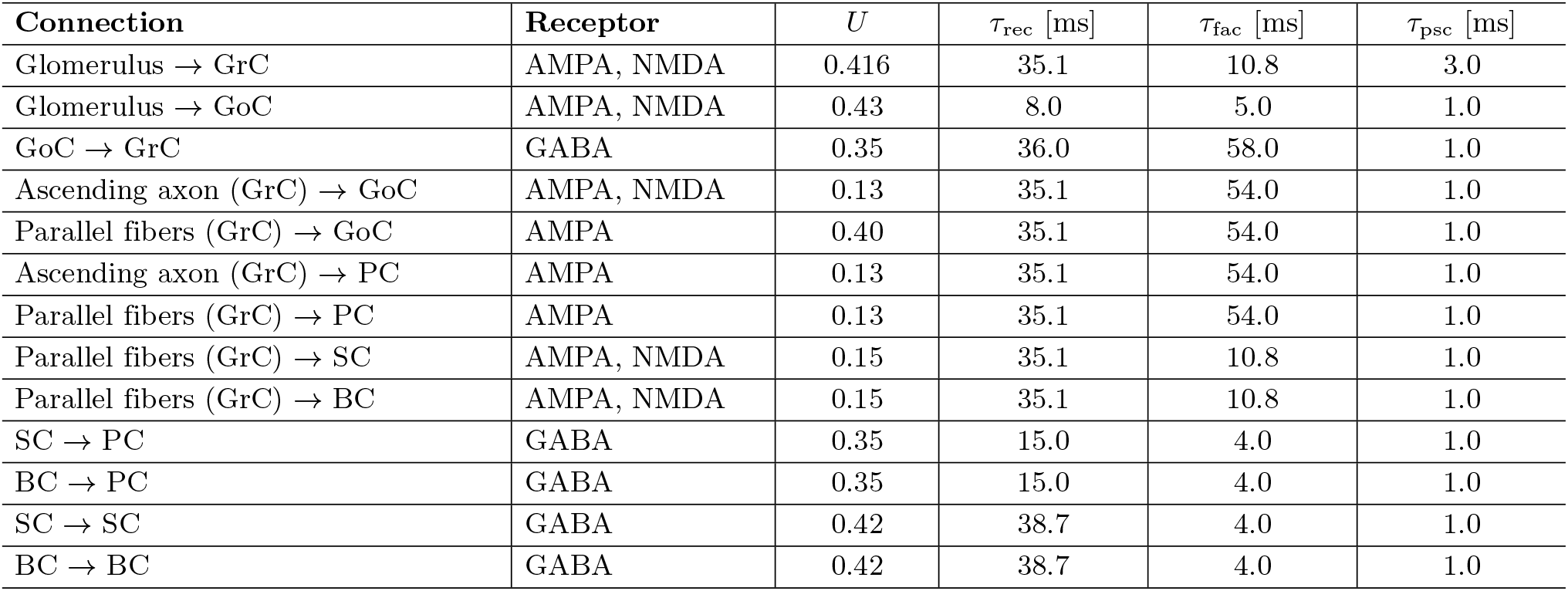
Short-term plasticity parameters for the cerebellar network synaptic connections. Connection-specific STP parameter sets were adopted from Masoli et al., 2022 [42].

| Connection | Receptor | $U$ | $\tau_{\text{rec}}$ [ms] | $\tau_{\text{fac}}$ [ms] | $\tau_{\text{psc}}$ [ms] |
| --- | --- | --- | --- | --- | --- |
| Glomerulus → GrC | AMPA, NMDA | 0.416 | 35.1 | 10.8 | 3.0 |
| Glomerulus → GoC | AMPA, NMDA | 0.43 | 8.0 | 5.0 | 1.0 |
| GoC → GrC | GABA | 0.35 | 36.0 | 58.0 | 1.0 |
| Ascending axon (GrC) → GoC | AMPA, NMDA | 0.13 | 35.1 | 54.0 | 1.0 |
| Parallel fibers (GrC) → GoC | AMPA | 0.40 | 35.1 | 54.0 | 1.0 |
| Ascending axon (GrC) → PC | AMPA | 0.13 | 35.1 | 54.0 | 1.0 |
| Parallel fibers (GrC) → PC | AMPA | 0.13 | 35.1 | 54.0 | 1.0 |
| Parallel fibers (GrC) → SC | AMPA, NMDA | 0.15 | 35.1 | 10.8 | 1.0 |
| Parallel fibers (GrC) → BC | AMPA, NMDA | 0.15 | 35.1 | 10.8 | 1.0 |
| SC → PC | GABA | 0.35 | 15.0 | 4.0 | 1.0 |
| BC → PC | GABA | 0.35 | 15.0 | 4.0 | 1.0 |
| SC → SC | GABA | 0.42 | 38.7 | 4.0 | 1.0 |
| BC → BC | GABA | 0.42 | 38.7 | 4.0 | 1.0 |

## Acknowledgments

This research has received funding from the European Union’s Research and Innovation Program Horizon Europe under the grant agreements No. 101147319 (EBRAINS 2.0 Project) to ED and CC and No 101137289 (Virtual Brain Twin Project) to DR, from the project PRIN 2022 (funded by the Italian Ministry of University and Research) 20228B2HN5 “cerebellarNEuromodulation in ATaxia: digital cerebellar twin to predict the MOVEment rescue (NEAT-MOVE)”to MDG and CC, and from NEXTGENERATION EU project MNESYS (PE0000006) (funded by the Ministry of University and Research, under National Recovery and Resilience Plan) “a Multiscale integrated approach to the study of the nervous system in health and disease”to DB and ED.

## References

1. De Zeeuw CI, Ten Brinke MM. Motor learning and the cerebellum. Cold Spring Harbor perspectives in biology. 2015;7(9):a021683.

2. Ciapponi C, Li Y, Osorio Becerra DA, Rodarie D, Casellato C, Mapelli L, et al. Variations on the theme: focus on cerebellum and emotional processing. Frontiers in systems neuroscience. 2023;17:1185752.

3. Douglas RJ, Martin KA. Canonical cortical circuits. Handbook of brain microcircuits. 2010:15–21.

4. Casanova MF. Canonical circuits of the cerebral cortex as enablers of neuroprosthetics. Frontiers Media SA;2013.

5. De Schepper R, Geminiani A, Masoli S, Rizza MF, Antonietti A, Casellato C, et al. Model simulations unveil the structure-function-dynamics relationship of the cerebellar cortical microcircuit. Communications biology. 2022;5(1):1240.

6. Geminiani A, Casellato C, Boele HJ, Pedrocchi A, De Zeeuw CI, D’Angelo E. Mesoscale simulations predict the role of synergistic cerebellar plasticity during classical eyeblink conditioning. PLOS Computational Biology. 2024;20(4):e1011277.

7. D’Angelo E, Antonietti A, Geminiani A, Gambosi B, Alessandro C, Buttarazzi E, et al. Linking cellular-level phenomena to brain architecture: the case of spiking cerebellar controllers. Neural Networks. 2025;188:107538.

8. Lorenzi RM, Geminiani A, Zerlaut Y, De Grazia M, Destexhe A, Gandini Wheeler-Kingshott CA, et al. A multi-layer mean-field model of the cerebellum embedding microstructure and population-specific dynamics. PLOS Computational Biology. 2023;19(9):e1011434.

9. Lorenzi RM, De Grazia M, Gandini Wheeler-Kingshott CAM, Palesi F, D’Angelo EU, Casellato C. Automated derivation of mean field models from spiking neural networks for the simulation of brain dynamics. bioRxiv. 2026:2026–03.

10. Masoli S, Solinas S, D’Angelo E. Action potential processing in a detailed Purkinje cell model reveals a critical role for axonal compartmentalization. Frontiers in Cellular Neuroscience. 2015;9:47.

11. Rizza MF, Locatelli F, Masoli S, Sánchez-Ponce D, Muñoz A, Prestori F, et al. Stellate cell computational modeling predicts signal filtering in the molecular layer circuit of cerebellum. Scientific Reports. 2021;11(1):3873.

12. Masoli S, Ottaviani A, Casali S, D’Angelo E. Cerebellar Golgi cell models predict dendritic processing and mechanisms of synaptic plasticity. PLoS computational biology. 2020;16(12):e1007937.

13. Masoli S, Tognolina M, Laforenza U, Moccia F, D’Angelo E. Parameter tuning differentiates granule cell subtypes enriching transmission properties at the cerebellum input stage. Communications biology. 2020;3(1):222.

14. Masoli S, Rizza MF, Soda T, Sánchez-Ponce D, Munoz A, Prestori F, et al. Cerebellar basket cell filtering of Purkinje cell responses elicited by low frequency parallel fibre transmission. Scientific Reports. 2025;15(1):25192.

15. Brunel N, Hakim V, Isope P, Nadal JP, Barbour B. Optimal information storage and the distribution of synaptic weights: perceptron versus Purkinje cell. Neuron. 2004;43(5):745–57.

16. Gagliano G, Monteverdi A, Casali S, Laforenza U, Gandini Wheeler-Kingshott CA, D’Angelo E, et al. Non-linear frequency dependence of neurovascular coupling in the cerebellar cortex implies vasodilation–vasoconstriction competition. Cells. 2022;11(6):1047.

17. Monteverdi A, Di Domenico D, D’Angelo E, Mapelli L. Anisotropy and frequency dependence of signal propagation in the cerebellar circuit revealed by high-density multielectrode array recordings. Biomedicines. 2023;11(5):1475.

18. Mapelli J, Gandolfi D, D’Angelo E. High-pass filtering and dynamic gain regulation enhance vertical bursts transmission along the mossy fiber pathway of cerebellum. Frontiers in cellular neuroscience. 2010;4:926.

19. Brain Scaffold Builder Developers. Brain Scaffold Builder (BSB);2024. Accessed March 2026. https://bsb.readthedocs.io/en/latest/.

20. Hines ML, Carnevale NT. The NEURON simulation environment. Neural Computation. 1997;9(6):1179–209.

21. Geminiani A, Casellato C, D’Angelo E, Pedrocchi A. Complex electroresponsive dynamics in olivocerebellar neurons represented with extended-generalized leaky integrate and fire models. Frontiers in Computational Neuroscience. 2019;13:35.

22. Brette R, Gerstner W. Adaptive exponential integrate-and-fire model as an effective description of neuronal activity. Journal of neurophysiology. 2005;94(5):3637–42.

23. Izhikevich EM. Which model to use for cortical spiking neurons? IEEE transactions on neural networks. 2004;15(5):1063–70.

24. Bahl A, Stemmler MB, Herz AV, Roth A. Automated optimization of a reduced layer 5 pyramidal cell model based on experimental data. Journal of neuroscience methods. 2012;210(1):22–34.

25. Pozzorini C, Mensi S, Hagens O, Naud R, Koch C, Gerstner W. Automated high-throughput characterization of single neurons by means of simplified spiking models. PLoS computational biology. 2015;11(6):e1004275.

26. Teeter C, Iyer R, Menon V, Gouwens N, Feng D, Berg J, et al. Generalized leaky integrate-and-fire models classify multiple neuron types. Nature communications. 2018;9(1):709.

27. Geminiani A, Casellato C, Locatelli F, Prestori F, Pedrocchi A, D’Angelo E. Complex dynamics in simplified neuronal models: reproducing Golgi cell electroresponsiveness. Frontiers in neuroinformatics. 2018;12:88.

28. Fortin FA, De Rainville FM, Gardner MAG, Parizeau M, Gagné C. DEAP: Evolutionary algorithms made easy. The Journal of Machine Learning Research. 2012;13(1):2171–5.

29. Deb K, Pratap A, Agarwal S, Meyarivan T. A fast and elitist multiobjective genetic algorithm: NSGA-II. IEEE transactions on evolutionary computation. 2002;6(2):182–97.

30. Van Geit W, De Schutter E, Achard P. Automated neuron model optimization techniques: a review. Biological cybernetics. 2008;99(4):241–51.

31. Druckmann S, Banitt Y, Gidon AA, Schürmann F, Markram H, Segev I. A novel multiple objective optimization framework for constraining conductance-based neuron models by experimental data. Frontiers in neuroscience. 2007;1:56.

32. Miettinen K. Nonlinear multiobjective optimization. vol. 12. Springer Science &Business Media;1999.

33. Wang XJ. Synaptic basis of cortical persistent activity: the importance of NMDA receptors to working memory. Journal of Neuroscience. 1999;19(21):9587–603.

34. Brunel N, Wang XJ. Effects of neuromodulation in a cortical network model of object working memory dominated by recurrent inhibition. Journal of computational neuroscience. 2001;11(1):63–85.

35. Kuhn A, Aertsen A, Rotter S. Neuronal integration of synaptic input in the fluctuation-driven regime. Journal of Neuroscience. 2004;24(10):2345–56.

36. Bernander O, Douglas RJ, Martin K, Koch C. Synaptic background activity influences spatiotemporal integration in single pyramidal cells. Proceedings of the National Academy of Sciences. 1991;88(24):11569–73.

37. Hill S, Tononi G. Modeling sleep and wakefulness in the thalamocortical system. Journal of neurophysiology. 2005;93(3):1671–98.

38. Plesser HE, Diesmann M, Gewaltig MO, Morrison A. NEST: the neural simulation tool. In: Encyclopedia of computational neuroscience. Springer;2022. p. 2187–9.

39. Plotnikov D, Rumpe B, Blundell I, Ippen T, Eppler JM, Morrison A. NESTML: a modeling language for spiking neurons. arXiv preprint arXiv:160602882. 2016.

40. DBBS Lab. Cerebellar Models;2024. GitHub repository, accessed: 2026-03-05. https://github.com/dbbs-lab/cerebellar-models.

41. Blank J, Deb K. Pymoo: Multi-objective optimization in python. Ieee access. 2020;8:89497–509.

42. Masoli S, Rizza MF, Tognolina M, Prestori F, D’Angelo E. Computational models of neurotransmission at cerebellar synapses unveil the impact on network computation. Frontiers in Computational Neuroscience. 2022;16:1006989.

43. Tsodyks M, Pawelzik K, Markram H. Neural networks with dynamic synapses. Neural computation. 1998;10(4):821–35.

44. Jiang HJ, Qi G, Duarte R, Feldmeyer D, van Albada SJ. A layered microcircuit model of somatosensory cortex with three interneuron types and cell-type-specific short-term plasticity. Cerebral Cortex. 2024;34(9):bhae378.

45. Mapelli J, D’Angelo E. The spatial organization of long-term synaptic plasticity at the input stage of cerebellum. Journal of Neuroscience. 2007;27(6):1285–96.

46. Rancz EA, Ishikawa T, Duguid I, Chadderton P, Mahon S, Häusser M. High-fidelity transmission of sensory information by single cerebellar mossy fibre boutons. Nature. 2007;450(7173):1245–8.

47. Rössert C, Pozzorini C, Chindemi G, Davison AP, Eroe C, King J, et al. Automated point-neuron simplification of data-driven microcircuit models. arXiv preprint arXiv:160400087. 2016.

48. Einevoll GT, Kayser C, Logothetis NK, Panzeri S. Modelling and analysis of local field potentials for studying the function of cortical circuits. Nature Reviews Neuroscience. 2013;14(11):770–85.

49. Mazzoni A, Lindén H, Cuntz H, Lansner A, Panzeri S, Einevoll GT. Computing the local field potential (LFP) from integrate-and-fire network models. PLoS computational biology. 2015;11(12):e1004584.

50. D’Angelo E, Rossi P, Taglietti V. Different proportions of N-methyl-D-aspartate and non-N-methyl-D-aspartate receptor currents at the mossy fibre-granule cell synapse of developing rat cerebellum. Neuroscience. 1993;53(1):121–30.

51. Zhou H, Lin Z, Voges K, Ju C, Gao Z, Bosman LWJ, et al. Cerebellar modules operate at different frequencies. eLife. 2014;3:e02536. doi:10.7554/eLife.02536.

52. Arancillo M, White JJ, Lin T, Stay TL, Sillitoe RV. In vivo analysis of Purkinje cell firing properties during postnatal mouse development. Journal of neurophysiology. 2015;113(2):578–91.

53. van Beugen BJ, Gao Z, Boele HJ, Hoebeek F, De Zeeuw CI. High frequency burst firing of granule cells ensures transmission at the parallel fiber to purkinje cell synapse at the cost of temporal coding. Frontiers in neural circuits. 2013;7:95.

54. Person AL, Raman IM. Purkinje neuron synchrony elicits time-locked spiking in the cerebellar nuclei. Nature. 2012;481(7382):502–5.

55. Marr D. Simple memory: a theory for archicortex. Philosophical Transactions of the Royal Society of London B, Biological Sciences. 1971;262(841):23–81.

56. Albus JS. A theory of cerebellar function. Mathematical biosciences. 1971;10(1-2):25–61.

57. Dean P, Porrill J, Ekerot CF, Jörntell H. The cerebellar microcircuit as an adaptive filter: experimental and computational evidence. Nature Reviews Neuroscience. 2010;11(1):30–43.

58. Solinas S, Nieus T, D’Angelo E. A realistic large-scale model of the cerebellum granular layer predicts circuit spatio-temporal filtering properties. Frontiers in cellular neuroscience. 2010;4:903.

59. Gurnani H, Silver RA. Multidimensional population activity in an electrically coupled inhibitory circuit in the cerebellar cortex. Neuron. 2021;109(10):1739–53.

60. Rössert C, Solinas S, D’Angelo E, Dean P, Porrill J. Model cerebellar granule cells can faithfully transmit modulated firing rate signals. Frontiers in cellular neuroscience. 2014;8:304.

61. Tirado MP, Ortigosa EM, Ros E, Garrido JA. A computational model of the cerebellar granular layer calibrated to experimental data for studying inhibition and sensory encoding. Scientific Reports. 2025;15(1):41788.

62. Yamazaki T, Tanaka S. A spiking network model for passage-of-time representation in the cerebellum. European Journal of Neuroscience. 2007;26(8):2279–92.

63. Lennon W, Hecht-Nielsen R, Yamazaki T. A spiking network model of cerebellar Purkinje cells and molecular layer interneurons exhibiting irregular firing. Frontiers in computational neuroscience. 2014;8:157.

64. Nieus T, Sola E, Mapelli J, Saftenku E, Rossi P, D’Angelo E. LTP regulates burst initiation and frequency at mossy fiber–granule cell synapses of rat cerebellum: experimental observations and theoretical predictions. Journal of neurophysiology. 2006;95(2):686–99.

65. D’Angelo E, De Filippi G, Rossi P, Taglietti V. Synaptic excitation of individual rat cerebellar granule cells in situ: evidence for the role of NMDA receptors. The Journal of physiology. 1995;484(2):397–413.

66. Kanichay RT, Silver RA. Synaptic and cellular properties of the feedforward inhibitory circuit within the input layer of the cerebellar cortex. Journal of Neuroscience. 2008;28(36):8955–67.

67. Segev I, London M. Untangling dendrites with quantitative models. Science. 2000;290(5492):744–50.

68. Parasuram H, Nair B, Naldi G, D’Angelo E, Diwakar S. A modeling based study on the origin and nature of evoked post-synaptic local field potentials in granular layer. Journal of Physiology-Paris. 2011;105(1-3):71–82.

69. Mapelli L, Di Domenico D, Sciacca G, Mainardi F, Ottaviani A, Monteverdi A, et al. Enhanced electrophysiological recordings in acute brain slices, spheroids, and organoids using 3D high-density multielectrode arrays. Plos one. 2025;20(9):e0328903.

70. Person AL, Raman IM. Synchrony and neural coding in cerebellar circuits. Frontiers in neural circuits. 2012;6:97.

71. Faris P, Pischedda D, Palesi F, D’Angelo E. New clues for the role of cerebellum in schizophrenia and the associated cognitive impairment. Frontiers in cellular neuroscience. 2024;18:1386583.

72. Vergani AA, Faris P, De Grazia M, Fusar-Poli P, Casellato C, D’Angelo EU. Cerebellar Degeneration Induces Timeliness Disruption, Paradoxical Information Flow and Network Overdrive in a Digital Model of Schizophrenia. Zenodo. 2026;Version 1.0. Available from: https://doi.org/10.5281/zenodo.18813588. doi:10.5281/zenodo.18813588.

73. Soda T, Mapelli L, Locatelli F, Botta L, Goldfarb M, Prestori F, et al. Hyperexcitability and hyperplasticity disrupt cerebellar signal transfer in the IB2 KO mouse model of autism. Journal of Neuroscience. 2019;39(13):2383–97.

74. Casellato C, Antonietti A, Garrido JA, Ferrigno G, D’Angelo E, Pedrocchi A. Distributed cerebellar plasticity implements generalized multiple-scale memory components in real-robot sensorimotor tasks. Frontiers in Computational Neuroscience. 2015;9:24.

75. Sgritta M, Locatelli F, Soda T, Prestori F, D’Angelo EU. Hebbian spike-timing dependent plasticity at the cerebellar input stage. Journal of Neuroscience. 2017;37(11):2809–23.

76. Ito S, Haufler D, Fraile JG, Dai K, Aman J, Chen G, et al. Deep-learning-assisted simulation of a cortical circuit: integrating anatomy, physiology and function. bioRxiv. 2026:2026–03.

77. D’Angelo E, Jirsa V. The quest for multiscale brain modeling. Trends in neurosciences. 2022;45(10):777–90.

